# ATML1-GIR1-TPL/TPR transcriptional repression module controls glucosinolates and giant cells in *Arabidopsis thaliana* sepals

**DOI:** 10.64898/2026.06.03.724713

**Authors:** Lauren E. Apprill, Bilal Ahmad, Aytug Ulutas, Amanda Agosto Ramos, Sumin Na, Samira R. Laytimi, Ayianna K. Bailey, Adara L. Warner, Titus R. Neumann, Young-Jin Lee, Brandon L. Garcia, Daniel J. Kliebenstein, Kathrin Schrick

## Abstract

Glucosinolates (GSLs) are sulfur- and nitrogen-containing secondary metabolites that serve as defense compounds in Arabidopsis and other members of the Brassicales. Although the enzymatic pathway that produces GSLs is well-studied, the upstream mechanisms that control their tissue-specific synthesis are poorly understood. We identified a novel repression module that transcriptionally regulates GSL levels in sepals, the modified leaves that protect reproductive tissues within the floral bud. GLABRA2 (GL2) INTERACTING REPRESSOR1 (GIR1) interacts directly with *Arabidopsis thaliana* MERISTEM LAYER1 (ATML1), an HD-Zip IV transcription factor known to be required for giant cell formation in the sepal epidermis. This interaction requires a predicted Zn finger of GIR1 and the C-terminal START adjacent domain (STAD) of ATML1. The *gir1* loss-of-function mutants exhibit excess giant cells, in contrast to *atml1* mutants which display fewer giant cells, supporting the role of GIR1 as a negative regulator of ATML1. We confirmed that GIR1 interacts with TOPLESS (TPL) and TOPLESS-RELATED (TPR) corepressors, and coimmunoprecipitation demonstrated that GIR1 acts as an adaptor protein connecting ATML1 and TPL/TPR. RNA sequencing revealed that numerous genes involved in GSL biosynthesis, including the key transcriptional regulator MYB29, are upregulated in *gir1* mutants. Consistent with the transcriptomic data, chemical analysis revealed that *gir1* mutants display elevated GSL levels in sepals. Mass spectrometry imaging confirmed high GSL accumulation in *gir1* sepals compared to wild type and *atml1*. Overall, our findings uncover a previously unrecognized link between cell expansion and GSL metabolism, suggesting strategies for engineering plants with cell-type specific GSL profiles.

**Significance Statement:** Plants belonging to the order Brassicales produce sulfur-containing glucosinolate (GSL) metabolites that serve in defense against herbivory. In cruciferous vegetables such as broccoli and kale, these compounds contribute to their unique flavors and health-promoting attributes. In agriculturally important oilseed crops, they affect the palatability of animal feeds. Here, we identified a novel transcriptional regulatory module that controls GSL biosynthesis in the epidermis of the sepal, the floral organ that protects the reproductive tissues. This regulatory module also controls cell expansion of specific cell types in the sepal, demonstrating a surprising connection between cell growth and a chemical defense pathway in plants. Our results suggest strategies for engineering crops with tissue-specific GSL profiles to fit agronomic needs.

## Introduction

Glucosinolates (GSLs) are secondary metabolites that naturally occur in the plant model *Arabidopsis thaliana* (hereafter Arabidopsis) and other members of the Brassicales including agriculturally important oilseed crops such as canola, as well as cruciferous vegetables such as broccoli, kale, and cabbage ^1^. On the plant side, GSLs serve as important defense compounds against herbivores and pathogens ^2,3^. Relevant to the human diet, GSLs contribute to the pungent flavors and health-promoting properties of cruciferous vegetables ^4,5^. Epidemiological studies have identified GSL byproducts, such as isothiocyanates, as bioactive derivatives possessing anti-inflammatory, antioxidant and anticarcinogenic properties ^6–9^. The hydrolysis of GSLs via myrosinase occurs when plant cells are damaged ^10,11^, or through catabolic breakdown by gut microbiota ^12,13^.

In Arabidopsis, GSL biosynthesis follows amino acid chain elongation from Met for aliphatic GSLs and from Trp for indolic GSLs. A series of enzyme reactions convert the amino acid derivatives to form the core GSL scaffold. Structural diversity arises from side-chain modifications catalyzed by hydroxylases, methyltransferases, and oxygenases ^14,15^. Overall pathway architecture is conserved across the Brassicales with lineage-specific expansions and neo-functionalizations determining biological activities of specific GSLs ^5,16,17^. Transcriptional regulation of the pathway is primarily governed by R2R3-MYB transcription factors. MYB28, MYB29, and MYB76 control genes of the aliphatic GSL pathway, while MYB34, MYB51, and MYB122 regulate the indolic pathway ^18–20^. Environmental and developmental cues are integrated in response to jasmonic acid, and interactions with bHLH and MYC transcription factors modulate the pathway under stress ^21^.

While the GSL biosynthesis pathway and its transcriptional control are extensively studied ^4^, the epidermal cell-type regulation and upstream molecular mechanisms that negatively control this pathway are poorly understood. GSL biosynthesis is thought to mainly occur in the vasculature ^22^, where GTR transporters mediate redistribution to specialized sulfur-rich storage cells in the phloem ^23,24^. Additionally, there is evidence for nonrandom distribution of GSLs in the leaf epidermis ^25^. *Arabidopsis thaliana* MERISTEM 1 (ATML1) is a member of the homeodomain leucine-zipper (HD-Zip) IV transcription factor family that is expressed in the outermost layer (L1) of vegetative and floral organs and is critical for formation of the epidermis ^26^. It functions redundantly with its paralog PROTODERMAL FACTOR2 (PDF2) to positively regulate the expression of L1-specific genes during epidermal differentiation ^27,28^. In Arabidopsis sepals, ATML1 overexpression promotes the development of giant cells that nearly cover the entire sepal whereas *atml1* loss-of-function results in defective giant cell formation ^29,30^.

In the present study, we identify a novel transcriptional repressor of the GSL biosynthesis pathway that regulates ATML1 activity. GIR1 (GLABRA2 (GL2)-interacting repressor) ^31^ contains an ethylene-responsive element binding factor-associated amphiphilic repression (EAR) motif implicated in TOPLESS (TPL) corepressor recruitment ^32^. Our genetic analysis revealed that GIR1 controls giant cell formation in an opposing manner to ATML1. We demonstrate that GIR1 directly interacts with ATML1 to modulate its activity at the post-translational level. GIR1 contains Cys motifs that are predicted to form a C4 Zn finger required for interaction with ATML1 through its START adjacent domain (STAD), a C-terminal domain of unknown function. Protein-protein interaction studies confirmed that GIR1 interacts with TPL as well as TOPLESS-RELATED (TPR) proteins through its EAR motif. We propose a model in which GIR1 engages the ATML1 transcription factor and TPL/TPR corepressor simultaneously to facilitate transcriptional repression of GSL pathway genes in a cell-type specific manner.

## Results

### The *gir1* mutants display excess giant cells on sepals

*GIR1* and *GIR1*-like genes encode small proteins of ∼100 amino acids that are thought to function in transcriptional repression ^31,32^, but whose molecular and biological activities are not clearly defined. Tomato and tobacco *GIR1* homologs have roles in trichome development ^33–35^. In cotton, *GIR1*-related members are implicated in fiber elongation ^36,37^. While performing genetic crosses with *gir1* mutants from Arabidopsis, we discovered an overproduction of giant cells on their sepals (**Fig. 1A,B**). Giant cells are elongated epidermal cells, visible by their large size and outward bulging. Sepals, the outermost floral organs, protect the reproductive tissues from the environment and control flower opening ^38^. *GIR2*, a paralog of *GIR1*, shares high nucleotide and predicted amino acid similarity to *GIR1* ^31^. However, no obvious phenotypes were observed in sepals of *gir2* null mutants compared to the control (**Fig. S1, Fig. 1C**). We constructed *gir1;gir2* double mutants and found that their giant cell phenotypes were indistinguishable from that of *gir1* single mutants (**Fig. S1, Fig. 1C**). These data show that *GIR1* but not *GIR2* negatively regulates giant cell formation on sepals.

**Figure 1.**
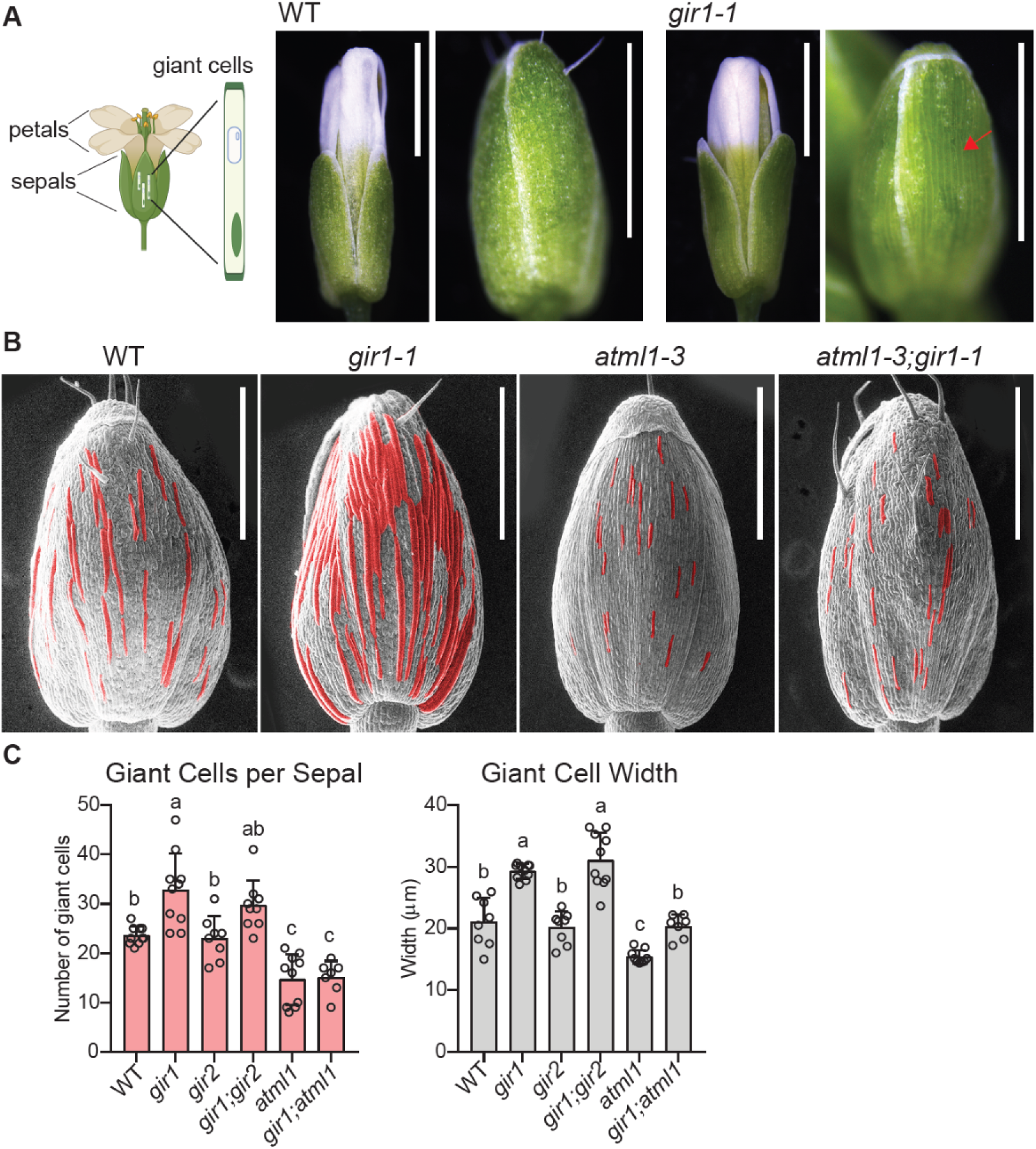
*gir1* mutant displays a giant cell phenotype in sepals. **(A)** Illustration of Arabidopsis flower indicating the anatomical positions of sepals and giant cells. Stereo microscope images show wild-type (WT) flowers (stage 14) and buds (stage 12) in comparison to *gir1* mutants, which display an increase in giant cell formation (arrow). Bar = 1 mm. **(B)** Comparison of giant cell phenotype on sepals from *gir1* versus *atml1* mutant. Enhanced giant cell formation is suppressed in the *gir1;atml1* double mutant, suggesting that *atml1* is epistatic to *gir1*. Scanning electron microscopy (SEM) images of stage 12 flowers with giant cells falsely colored red. Bar = 500 μm. **(C)** Quantification of giant cell numbers per sepal and giant cell width for WT in comparison to *gir1, gir2, gir1;gir2, atml1*, and *gir1;atml1.* SD are indicated for n=7-10 biological replicates. Significant differences were assessed by one-way ANOVA, Tukey’s test, and are indicated by letters (p value < 0.05). Images of *gir2* and *gir1;gir2* mutant buds are shown in **Fig. S1**.

### Giant cell phenotypes reveal genetic interaction between *GIR1* and *ATML1*

The *atml1* loss-of-function mutants were previously reported to exhibit defective giant cell formation ^30^. The giant cell overproduction we observed in *gir1* mutants is similar to that reported in lines overexpressing *ATML1* ^29^. Thus, *GIR1* appears to function antagonistically to *ATML1*. To further test the relationship between the *gir1* and *atml1* mutations we performed genetic crosses. We found that *atml1* single mutants and *gir1;atml1* double mutants have similarly low numbers of giant cells, indicating that *atml1* is genetically epistatic to *gir1* (**Fig. 1B**). Quantification of giant cell numbers and giant cell widths revealed significant increases in both metrics in *gir1* mutants in comparison to wild type and *atml1*, whereas this phenotype was lost in the *gir1;atml1* double mutant (**Fig. 1C**). Thus, *gir1* excess giant cell formation depends on *ATML1* activity. These results suggest that the balance between levels of *ATML1* and *GIR1* activities is important in determining giant cell fate within the sepal.

### GIR1 complements the excess giant cell phenotype of *gir1* mutants

Next, a complementation experiment was conducted to verify that the giant cell phenotype is due to the loss of *GIR1* function. We generated *gir1* transgenic lines carrying a genomic fragment containing the native promoter and coding region of wild-type *GIR1* (**Fig. 2A, 2B**). Observation and quantification of giant cells versus small cells on sepals confirmed rescue of the *gir1* mutant phenotype (**Fig. 2C, 2D**). From sequence alignments and AlphaFold models ^39^, we identified conserved motifs which we targeted for mutational analysis (**Figs. 2A, 3A**). The EAR motif is a transcriptional repression motif consisting of LxLxL (x = any amino acid) ^40,41^ found in the GIR1 N-terminus. Two conserved Cys motifs (CPxC), located within the C-terminus of GIR1, are predicted to form a Zn finger comprised of four Cys residues.

**Figure 2.**
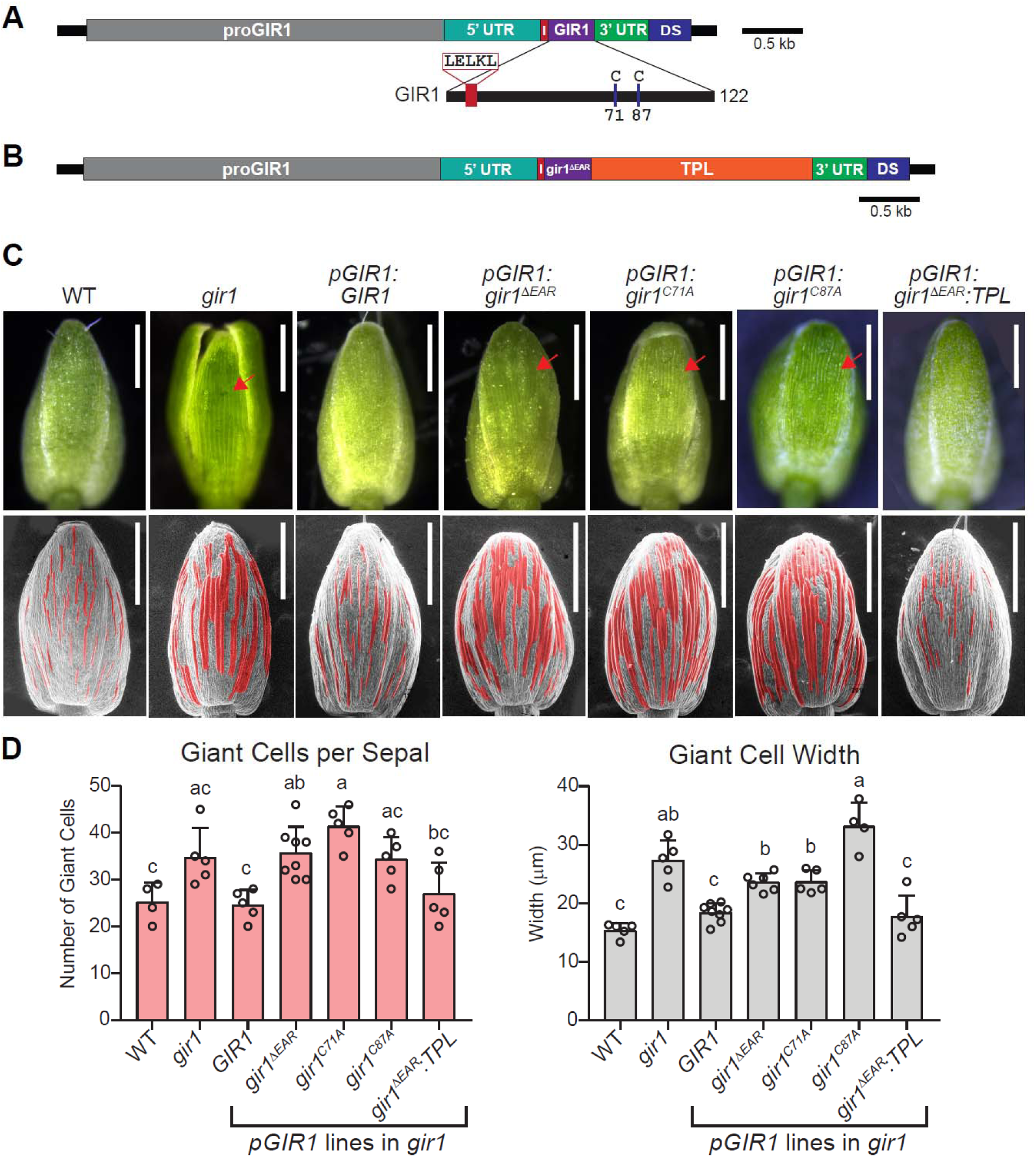
The EAR motif and Cys residues of the Zn finger are required for rescue of the giant cell phenotype by the *GIR1* native promoter construct. **(A) and (B)** Schematic of the transgenic native promoter constructs for *GIR1* wild-type and mutants affecting the function of the EAR motif and predicted Zn finger. **(A)** The *GIR1* native promoter (*pGIR1*) was used to drive expression of *GIR1*, and the *gir1^ΔEAR^* and *gir1^C68A^* mutants, and **(B)** expression of C-terminal fusion of TPL to *gir1^ΔEAR^*. **(C)** Stereo microscopy images of buds (stage 12) (top) and matching scanning electron microscopy images (bottom) with genotypes and constructs shown in **(A)** and **(B)** indicated. Giant cells are false-colored red. The *GIR1* and *gir1^ΔEAR^*-*TPL* constructs rescued *gir1* phenotype while *gir1* mutants (*gir1^ΔEAR^*, *gir1*^C71A^, and *gir1^C87A^*) failed to rescue. Bar = 500 μm. **(D)** Number of giant cells along with average giant cell width per sepal were quantified for the genotypes in **(C)**. SD are indicated for n=7-10 biological replicates. Significant differences were assessed by one-way ANOVA, Tukey’s test, and are indicated by letters (p value < 0.05).

**Figure 3.**
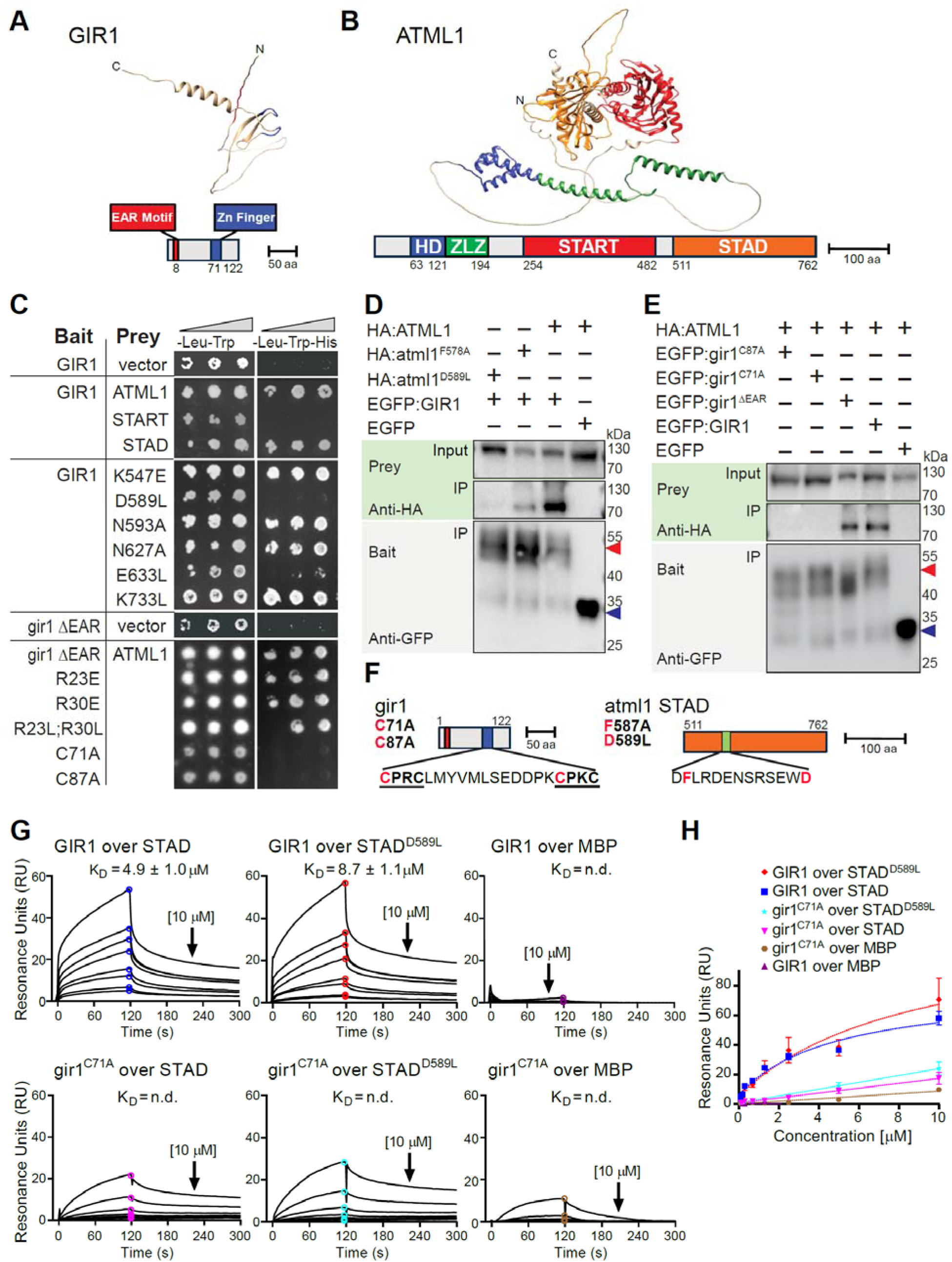
GIR1 interacts directly with HD-Zip IV transcription factor ATML1 via its Zn finger motif. **(A)** and **(B)** Predicted structures of GIR1 **(A)** and ATML1 **(B)** (AlphaFold; ^39^) visualized with UCSF ChimeraX ^58^. **(A)** GIR1 EAR motif is at the N-terminus of the protein while the Zn finger lies within the C-terminus. **(B)** ATML1 contains a HD, homeodomain; ZLZ, zipper–loop–zipper; START, steroidogenic acute regulatory (StAR) related lipid transfer domain; and STAD, START adjacent domain. **(C)** Y2H assays reveal that GIR1 interacts with ATML1 through STAD, and that STAD residues D589, N627, and E633 are critical for interaction. Cys residues (C71, C87) within the predicted Zn finger of GIR1 are important for interaction with ATML1, whereas the EAR motif and Arg residues within the N-terminus of GIR1 (R23, R30) are not required. The empty prey vector served as a negative control. Growth on - Leu-Trp signifies presence of bait and prey plasmids, while growth on selective media (-Leu-Trp-His) indicates protein interaction. Four-fold dilutions of yeast are indicated left to right. **(D)** and **(E)** Co-IP experiments confirmed GIR1-ATML1 interaction in planta using *Nicotiana benthamiana* transient expression. EGFP- and HA-tagged proteins served as baits and preys, respectively. Input indicates the initial protein amount for the Co-IP. Input and immunoprecipitated (IP) proteins were detected in western blots with anti-HA and anti-GFP antibodies. **(D)** ATML1 STAD residues F578 and D589 are important for physical interaction with GIR1. **(E)** The Cys mutants (gir1^C87A^, gir1^C71A^) failed to interact with ATML1 while the gir1^ΔEAR^ interaction with ATML1 was unaffected, indicating the importance of the GIR1 Zn finger for ATML1 interaction. **(F)** Schematic of mutations that impair the GIR1-ATML1 interaction. For GIR1, the C71A and C87A mutations affect the Zn finger motif. For ATML1 STAD, the F578A and D589L mutations affect aromatic and acidic residues, respectively, within an acidic patch. **(G)** and **(H)** SPR analysis of ATML1 STAD and atml1 STAD^D589L^ with GIR1 and gir1^C71A^ confirmed direct in vitro interaction between ATML1 STAD and GIR1. Purified MBP-tagged ATML1 STAD, atml1 STAD^D589L^, or MBP alone was immobilized on a SPR sensor chip. Purified GIR1 or gir1^C71A^ was injected in a series of 2-fold dilutions ranging from 0.08 – 10 µM. See **Fig. S4**. **(G)** Reference and background-corrected sensorgrams from a representative injection series are shown as black lines. Calculated steady-state equilibrium dissociation constants (K_D_) are shown as mean values from triplicate injections, and ± indicates SD. The data were fit to a steady-state model of binding. No fit or resulting K_D_ could be determined for the gir1^C71A^ mutant. See **Table S1**. **(H)** Steady-state affinity responses (circles) in **(G)** along with corresponding fits for each series are shown as concentration versus response plots (colored lines). Error bars represent the SD of three independent injections.

We utilized Cys to Ala mutagenesis to test whether the Zn finger is required for *GIR1* regulation of giant cell formation. Transgenic lines carrying the genomic fragment with the *gir1^C71A^* or *gir1^C87A^*mutations failed to complement the *gir1* giant cell phenotype (**Figs. 2A, 2C, 2D**). Similarly, *gir1*^Δ*EAR*^ transgenic lines carrying a genomic fragment harboring a deletion of the EAR motif failed to rescue the *gir1* giant cell phenotype (**Figs. 2A, 2C, 2D**). Since the EAR motif is required to recruit the TPL corepressor, we asked whether a translational fusion of TPL to the gir1^ΔEAR^ mutant protein restores function of GIR1. The TPL C-terminal 944 amino acid coding region was fused to the C-terminus of the *gir1*^Δ*EAR*^ genomic sequence (**Fig. 2B**). Phenotypic characterization showed that the *proGIR1:gir1*^Δ*EAR*^*:TPL* transgene complemented the giant cell phenotype, similar to the wild-type *pro:GIR1:GIR1* (**Figs. 2C, 2D**). This result indicates that GIR1 recruits TPL through the EAR motif.

Next, we repeated the complementation experiment with *gir1* mutants utilizing the cauliflower mosaic virus strong constitutive 35S promoter (**Fig. S2A**). *GIR1* overexpression similarly showed complementation of *gir1* giant cell phenotype, and quantification revealed a trend in the reduction of giant cell formation in the *35S:GIR1* lines (**Fig. S2B, S2C**). In contrast, 35S overexpression lines carrying the *gir1*^Δ*EAR*^, *gir1^C71A^* or *gir1^C87A^* mutations exhibited giant cell phenotypes indistinguishable from the *gir1* mutant (**Fig. S2B, S2C**). Thus, *GIR1* requires the EAR motif and the Cys residues within the predicted Zn finger to effectively suppress the giant cell overproduction in sepals.

### GIR1 protein directly interacts with ATML1 STAD

We utilized both in vivo and in vitro assays to test physical interaction between GIR1 and ATML1, considering the predicted structural features of each protein. Aside from the EAR motif and the predicted Zn finger, the GIR1 protein appears highly disordered (**Fig. 3A**). The ATML1 transcription factor contains a HD DNA binding domain and a leucine zipper dimerization domain, as well as the steroidogenic acute regulatory (StAR) protein-related lipid transfer (START) and STAD (**Fig. 3B**).

Using yeast 2-hybrid (Y2H) assays, we confirmed interaction between GIR1 and ATML1 and mapped the site of interaction to STAD for ATML1 (**Fig. 3C**). We utilized ATML1 as a prey in our Y2H assays since bait autoactivation assays showed that the ATML1 full-length protein displays autoactivation (**Fig. S3**). Among the structural features of interest, we noted that STAD contains a short acidic patch of 13 amino acids (**Fig. S3A**) and found that STAD, but not START, behaves as an autoactivation domain in yeast (**Fig. S3B**). Several missense mutations within STAD (K547E, F578A, D589L, E633L) resulted in loss of bait autoactivation. Our Y2H assays with STAD mutants confirmed the importance of D589, which is located within the STAD acidic patch, as well as D633, for interaction with GIR1 (**Fig. 3C**). We used Y2H assays to determine which features of GIR1 are critical for binding to ATML1. Mutants affecting the EAR motif (*gir1*^Δ*EAR*^) were proficient in interaction with ATML1, whereas the Cys (CPxC) motif mutants *gir1^C71A^*and *gir1^C87A^* failed to interact with ATML1 (**Fig. 3C**).

The GIR1-ATML1 protein interactions were validated by co-immunoprecipitation (Co-IP) using transient expression of HA- and GFP-tagged proteins in *N. benthamiana*. The EGFP:GIR1 bait protein strongly interacted with HA:ATML1 (**Figs. 3D, 3E**). Furthermore, specific mutations mapping to STAD reduced or abolished the interaction, consistent with the Y2H assays. The HA:atml1^F578A^ and HA:atml1^D589L^ mutants showed a weaker and no detectable interaction, respectively (**Fig. 3D**). Consistent with the Y2H results, the *gir1*^Δ*EAR*^ mutant was proficient in Co-IP with ATML1, while the Cys (CPxC) motif mutants *gir1^C71A^* and *gir1^C87A^* failed to interact with ATML1 (**Fig. 3E**). Taken together, the Co-IP experiments confirmed the critical importance of the Cys residues (C71, C87) of GIR1, and the STAD residues (F587, D589) of ATML1 for interaction (**Fig. 3F**).

To evaluate the interaction in a biochemical assay, we performed surface plasmon resonance (SPR) experiments with recombinant ATML1 STAD and GIR1 proteins purified from *E. coli* (**Fig. S4**). The SPR sensorgrams revealed binding between ATML1 STAD and GIR1 with a low micromolar equilibrium dissociation constant (K_D_ = 4.9 +/- 1.0 µM), while the atml1 STAD^D589L^ mutant showed nearly 2-fold lower affinity (K_D_ = 8.7 +/- 1.1 µM) (**Fig. 3G, 3H, Table S1**). In contrast, ATML1 STAD failed to bind effectively with the mutant gir1^C71A^ protein, and the maltose binding protein, used as a negative control, also failed to bind GIR1 and gir1^C71A^ (**Fig. 3G, 3H, Table S1**). The results establish a direct interaction between ATML1 STAD and GIR1 and further highlight the role of the predicted Zn finger of GIR1 in binding to STAD.

### ATML1, GIR1, and TPL/TPR proteins form a ternary complex

A previous study showed that GIR1 binds the TPL corepressor, suggesting its role as an adaptor protein ^32^. We conducted Co-IP experiments with EGFP-tagged GIR1 wild-type and mutant proteins and HA-tagged TPL to address the functional requirements for this interaction. Our results show that the EAR motif is necessary for TPL binding while the Cys (CPxC) motifs are dispensable for this interaction (**Fig. 4A**). Since TPL is a member of a small gene family, we used Co-IP to address whether several TPL-related (TPR) proteins also interact with GIR1. Our results show that four family members (TPR1, TPR2, TPR3, TPR4) bind to GIR1 in a manner similar to TPL (**Fig. 4B**). We validated the GIR1-TPL/TPR interaction results with Y2H assays and additionally confirmed that gir1^ΔEAR^ mutant fails to interact with these corepressors, although both mutant and wild-type display similar levels of protein expression (**Fig. 4C, Fig. S5**). Another corepressor protein, SAP18, failed to interact with GIR1, serving as a negative control, along with the empty vector (**Fig. 4C**).

**Figure 4.**
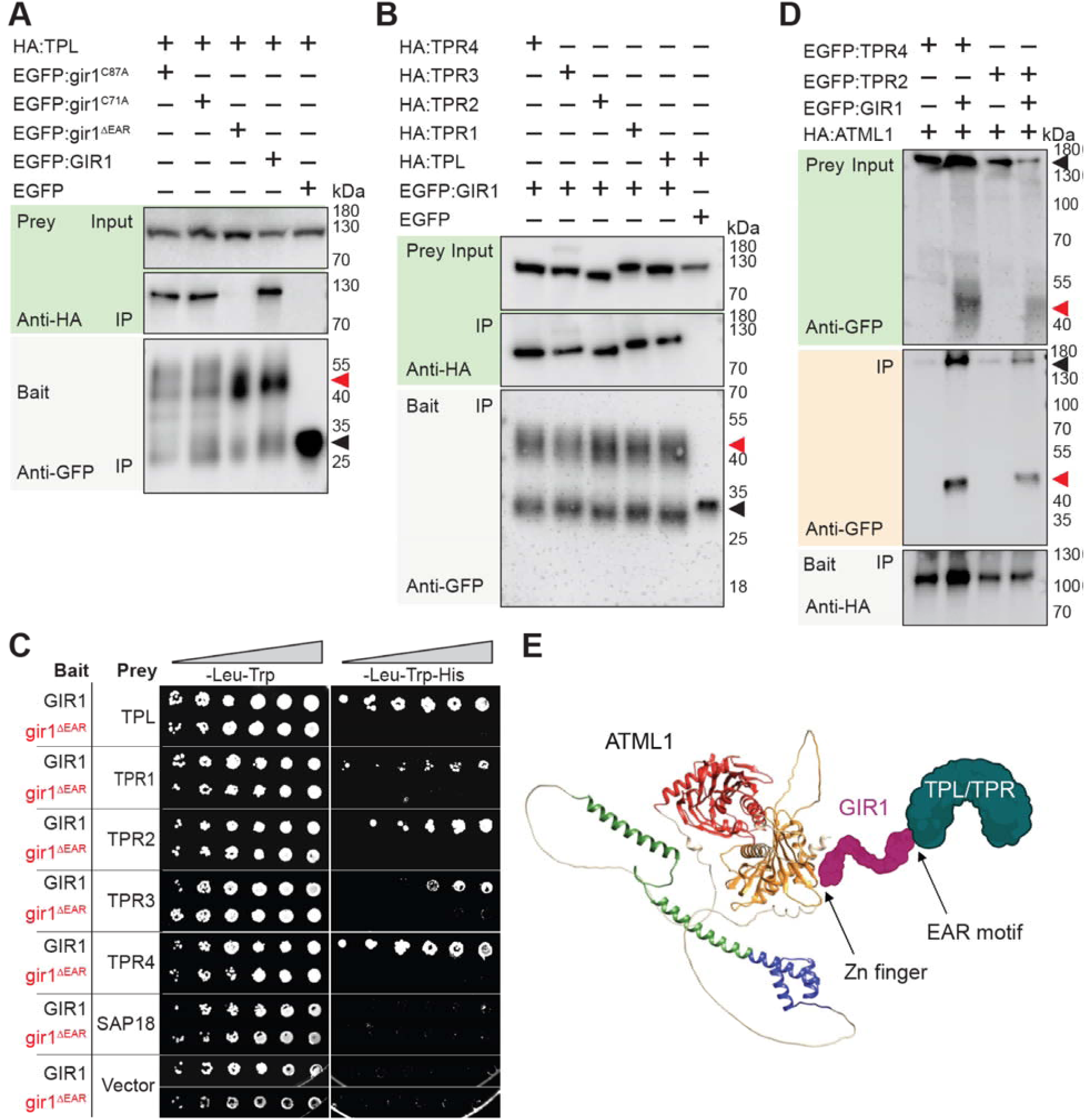
GIR1 interacts with TPL and TPRs via its EAR motif. **(A) and (B)** Co-IP experiments demonstrate GIR1-TPL and GIR1-TPR interactions in planta using *N. benthamiana* transient expression. EGFP- and HA-tagged proteins served as baits and preys, respectively. Input indicates the initial protein amount used for the Co-IP. Input and immunoprecipitated (IP) proteins were detected in western blots with anti-HA and anti-GFP antibodies. **(A)** Co-IP shows that the GIR1 EAR motif, but not its Zn-finger associated Cys residues, is required for GIR1-TPL interaction. EGFP-tagged GIR1, gir1 mutants (gir1^C71A^, gir1^C87A^, gir1^ΔEAR^) (red arrowhead), or EGFP control (black arrowhead) served as baits whereas HA-tagged TPL served as prey. **(B)** Co-IP shows GIR1 interaction with TPL and TPR proteins. EGFP-tagged GIR1 (red arrowhead) or EGFP negative control (black arrowhead) served as bait whereas HA-tagged TPL or TPR proteins served as preys. **(C)** Y2H assays confirm that GIR1 interacts with TPL and TPR proteins and that the GIR1 EAR motif is required for this interaction. SAP18 and empty vector served as negative controls. Growth on -Leu-Trp signifies presence of bait and prey plasmids, while growth on selective media (-Leu-Trp-His) indicates protein interaction. Four-fold dilutions of yeast are indicated left to right. **(D)** Co-IP demonstrates that GIR1 serves as a bridge connecting ATML1 and TPR proteins (TPR2, TPR4). HA-ATML1 and EGFP-TPR2 (or TPR4) were coexpressed with and without EGFP:GIR1 in *N. benthamiana*, and proteins were immunoprecipitated using anti-HA magnetic beads. Proteins were detected with anti-HA or anti-GFP antibodies in western blots. Black arrowheads indicate EGFP:TPR2 and EGFP:TPR4 prey proteins in the input (top blot) and co-immunoprecipitated proteins (middle blot) in the presence or absence of GIR1. Red arrowheads indicate EGFP:GIR1 in the input (top blot) and coimmunoprecipitated proteins (middle blot). The HA:ATML1 bait is shown in the bottom blot. **(E)** Model of the tripartite complex. ATML1 binds GIR1 via its C-terminal STAD segment, while GIR1 binds ATML1 via its Zn finger. The EAR motif of GIR1 is required for binding to TPL or TPR proteins. Predicted contact sites of the Zn finger and EAR motif are indicated by arrows.

Since GIR1 interacts with ATML1 (**Fig. 3**) and TPL/TPR, we investigated whether GIR1 can interact with both proteins simultaneously, thereby forming a tripartite complex. HA:ATML1 was co-expressed with EGFP-tagged TPR2 and TPR4 in *N. benthamiana* either in absence or presence of EGFP:GIR1. Immunoprecipitation with anti-HA magnetic beads showed that HA-ATML1 pulled down TRP2 and TPR4 in the presence of GIR1 (**Fig. 4D**). In contrast, little or no interaction was observed between ATML1 and TPR2/TPR4 when co-expressed in the absence of GIR1. Our results demonstrate that GIR1 facilitates the formation of a ternary complex through simultaneous interactions with ATML1 and TPL/TPR family members such as TPR2 and TPR4 (**Fig. 4E**).

### GSL pathway genes are upregulated in sepals from *gir1* mutants

To understand the molecular basis of the excess giant cell phenotype in *gir1* mutants, we performed RNA-seq on sepals from stage 12 buds of *gir1* and *atml1* mutants, and wild type (**Datasets S1-S7**). As expected, the transcript levels of *GIR1* and *ATML1* were decreased in the corresponding mutants in comparison to wild type (**Fig. 5A**). Strikingly, the *gir1* mutants exhibited more upregulated genes relative to wild type (**Fig. 5B**). In contrast, *atml1* mutants exhibited more downregulated genes. The upregulation of genes in the *gir1* mutant in comparison to both wild type and *atml1* (**Figs. 5C**, **5D**; **Fig. S6)**, provides support for the hypothesis that *GIR1* functions as a repressor of the giant cell fate, while *ATML1* functions as an activator of giant cell fate.

**Figure 5.**
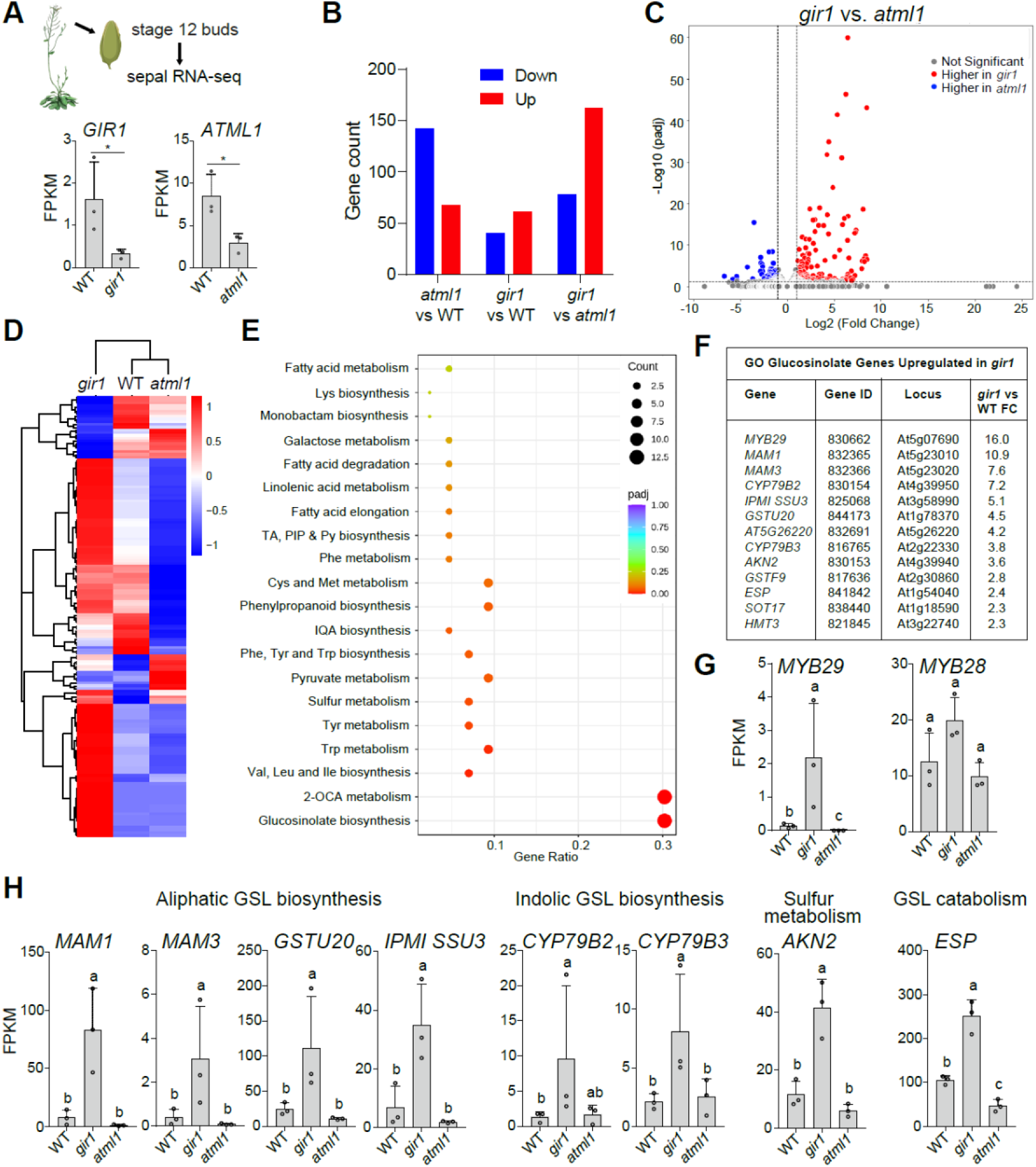
Differential expression analysis suggests that GIR1 acts as a repressor of ATML1 to control GSL biosynthesis genes in sepals. **(A)** Sepals from stage 12 buds were taken for RNA-seq analysis of wild type (WT), *gir1* and *atml1*. *GIR1* and *ATML1* mRNA levels are decreased in each corresponding mutant background. FPKM, fragments per kilobase per million mapped reads. Asterisk indicates significant difference to WT (unpaired *t*-test with Welch’s correction, p < 0.05) **(B)** Binary comparison of differentially expressed genes (DEGs) in sepals for WT, *gir1* mutant, and *atml1* mutant genotypes. Blue bars correspond to downregulated genes and red bars represent upregulated genes. **(C)** Volcano plot illustrates the greater number of upregulated genes in *gir1* in comparison to *atml1*. Red dots represent upregulated genes and blue dots represent down-regulated genes. Genes with unchanged expression are indicated by grey dots. **(D)** Heatmap shows overview of DEGs between *atml1*, *gir1*, and WT sepals. Averaged relative expression between n=3 biological replicates is graded from low (blue) to high (red) (P < 0.05). A dendrogram depicting clustering of gene expression patterns is shown on the left of the heatmap. Individual replicates are shown for each genotype in Fig. S6. **(E)** Dot plot visualization of gene ontology (GO) enrichment analysis of upregulated genes of *gir1* vs. *atml1*. The y-axis shows GO terms while the x-axis shows the proportion of total genes identified in the gene set which are also associated with the GO term. Dot color reflects the statistical significance (-log10 (padj-value)) whereas the dot size corresponds to the number of genes in each GO term. **(F)** List of upregulated genes in *gir1* sepals associated with the GO term GSL biosynthesis. Expression fold difference between *gir1* and WT is provided for each gene. **(G)** Sepal RNA-seq data for *MYB29* and *MYB28*, two redundant regulators of GSL biosynthesis. *MYB29* expression levels are >10-fold higher in *gir1* as compared to WT, while *MYB29* transcripts were not detected in *atml1*. **(H)** Sepal RNA-seq data for other GSL biosynthesis genes upregulated in *gir1* mutants, including aliphatic (*MAM1*, *MAM3*, *GSTU20*, *IPMI SSU3*) and indolic (*CYP79B2*, *CYP79B3*) GSL biosynthesis genes, as well as a genes involved in sulfur metabolism (*AKN2*) and GSL catabolism (*ESP*), respectively. In **(G)** and **(H)**, significant differences between genotypes are marked by letters (one-way ANOVA, Tukey’s test, p < 0.05). RNA-seq data for additional genes is shown in **Figs. S7-S8** and **Table S2**.

Gene ontology (GO) and overrepresentation analysis initially revealed the upregulation of 13 GSL pathway genes in *gir1* mutants (**Figs. 5E, 5F**). The upregulated genes included *MYB29*, a transcription factor gene implicated in aliphatic GSL production ^42^, and *MAM1*, an aliphatic GSL biosynthesis gene, both of which were upregulated >10-fold in *gir1* mutants (**Figs. 5F, 5G, 5H**). Genes significantly upregulated in *gir1* included other aliphatic GSL biosynthesis genes (*MAM3, GSTU20, IPMI SSU3*), as well as genes implicated in indolic GSL biosynthesis (*CYP79B2, CYP79B3*), sulfur metabolism (*AKN2*) and GSL catabolism (*ESP*) (**Fig. 5H**). The *MYB29* paralog, *MYB28*, also showed an upregulated trend in *gir1* mutants (**Fig. 5F**).

Additionally, the GO analysis revealed that genes related to 2-oxocarboxylic acid metabolism are highly enriched in *gir1* (**Fig. 5E**). Also referred to as 2-keto acid, 2-oxocarboxylic acid is a key intermediate in amino acid synthesis. Met undergoes transamination to produce 2-oxocarboxylic acid in the initial steps of aliphatic GSL biosynthesis ^43^. A series of reactions elongate the Met side chain prior to GSL core structure formation. Upregulated genes involved in 2-oxocarboxylic acid metabolism indicate an increase in elongated chain Met, consistent with the elevated production of aliphatic GSLs in *gir1*.

Further scanning of the RNA-seq data uncovered numerous other GSL pathway genes that were significantly upregulated or showed increased trends in *gir1* mutants in comparison to wild type and *atml1.* These include 21 aliphatic GSL genes (**Fig. S7, Table S2**) and 9 indolic GSL genes, two of which are also active in the aliphatic pathway (**Fig. S8, Table S2**). Benzenic GSL pathway genes were not differentially expressed in *gir1*, while one endoplasmic reticulum (ER) body GSL-related gene, *NAI1*, was significantly upregulated (**Fig. S8, Table S2**). Consistent with our hypothesis that *ATML1* is a positive regulator of GSL biosynthesis, *MYB29* and several other GSL-related genes were significantly downregulated or showed a decreased trend in *atml1* mutants. While we found evidence for *GIR1*-mediated transcriptional regulation of the aliphatic pathway through *MYB29* and to a lesser extent, *MYB28*, the regulators of the indolic pathway, *MYB34* and *MYB51*, showed no differential expression (**Fig. S8**), and transcript levels of *MYB76* and *MYB122* were not detected in the RNA-seq data (**Datasets S1-S7**).

### GSL levels are elevated in sepals from *gir1* mutants

To address whether upregulation of GSL biosynthesis pathway genes leads to changes in GSL production, we collected stage 12 sepals from wild type and from *gir1* and *atml1* mutants. To serve as a control, we also concurrently harvested bud tissues lacking sepals. Extracts were prepared for high performance liquid chromatography-diode-array detection (HPLC-DAD) (**Fig. 6A**). In wild-type and *atml1*, lower levels of total GSLs were detected in sepals in comparison to buds lacking sepals (**Fig. 6B, Table S3**). In contrast, *gir1* mutant sepals and buds lacking sepals contained similar levels of total GSLs in both tissues, due to a marked increase in GSLs in *gir1* sepals. Wild-type *GIR1*, when expressed under its native promoter, complemented the *gir1* GSL phenotype in sepals, while the mutant *gir1^C71A^* transgene did not rescue (**Fig. 6B-6D**). The results establish the importance of *GIR1* and its predicted Zn finger in controlling GSL levels in sepals.

**Figure 6.**
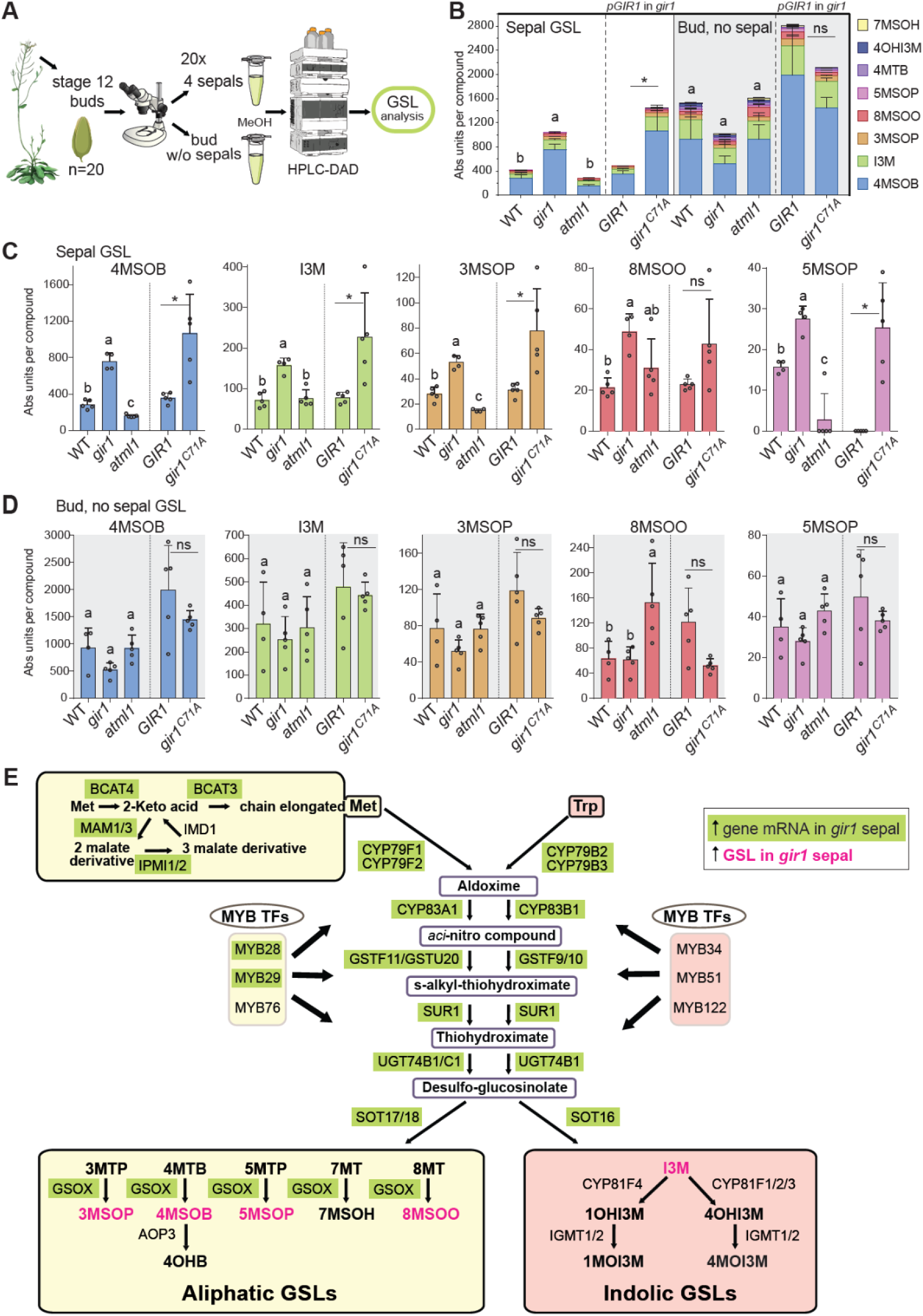
GSL levels are elevated in sepals from *gir1* mutants. **(A)** Schematic of workflow for GSL analysis. Stage 12 buds were dissected under the stereo microscope. Sepals were separated from the other bud tissues and transferred to 100% methanol prior to extraction, desulfation, and detection by HPLC-DAD. GSLs were quantified from absorbance peaks. **(B-D)** HPLC-DAD analysis uncovered elevated levels of GSLs in *gir1* mutant sepals in comparison to wild type (WT) and *atml1* (**Dataset S8**). Comparison of transgenic *gir1* lines expressing *pGIR1:GIR1* versus mutant *pGIR1:gir1^C71A^* constructs was performed in a separate experiment (**Dataset S9**). Wild-type *GIR1* but not the mutant *gir1^C71A^* rescued the *gir1* mutant phenotype in sepals. Full names of GSLs and information on their properties are given in **Table S4**. Significant differences between WT, *gir1* and *atml1* are marked by letters (one-way ANOVA, Tukey’s test, p < 0.05), and between transgenic lines by asterisk (unpaired *t*-test with Welch’s correction, p < 0.05); ns, not significant. **(B)** Stacked bar graphs illustrate GSL composition in sepals (left panel) in comparison to buds lacking sepals (right panel, grey shading). The key for the GSL compounds (right) is ordered from top to bottom from the least- to most-abundant compounds in sepals. 4MSOB is most abundant in both tissues, followed by I3M. The GSL profile in buds without sepals additionally includes the minor compounds 4MTB, 4OHI3M, and 7MSOH. **(C)** Bar graphs indicate that GSLs detected in sepals are elevated in *gir1* mutants, compared to WT, for all five compounds that were detected by HPLC-DAD (4MSOB, I3M, 3MSOP, 8MSOO, 5MSOP). The *atml1* mutants exhibit decreases in 3MSOP, 4MSOB, and 8MSOO. **(D)** Bar graphs show that buds without sepals failed to show consistent differences in GSL profiles between the genotypes. **Datasets S8-S9** show GSL profiles for the minor compounds. **(E)** Model of the GSL biosynthesis pathway in Arabidopsis. Aliphatic and indolic GSLs are derived from Met and Trp, respectively. The side-chain elongation of Met is shown at the top left. Core GSL scaffold formation is depicted in the center. The downstream enzymatic reactions specific to aliphatic and indolic GSLs are shown in the bottom left and right boxes, respectively. MYB transcription factors that control the aliphatic and indolic components are shown on the flanks of the core pathway. Genes upregulated in the *gir1* mutants are indicated in green boxes, and compounds detected by HPLC-DAD in sepals are shown in magenta. Modified from ^59^.

Overall, five GSLs (4MSOB, I3M, 3MSOP, 8MOSOO, 5MSOP) were detected in sepals and buds lacking sepals, and three additional minor GSLs (4MTB, 4OHI3M, 7MSOH) were detected in buds (**Datasets S8-S9**). Four aliphatic GSLs (4MSOB, 3MSOP, 8MOSOO, 5MSOP) and one indolic GSL (I3M) were significantly increased about two-fold in *gir1* sepals in comparison to wild-type sepals (**Fig. 6B, 6C, Table S3**). In contrast, *atml1* sepals exhibited a reduction in GSL levels for three of the aliphatic compounds (4MSOB, 3MSOP, 5MSOP). Notably, the quantitative differences in GSL levels were restricted to sepals. In buds lacking sepals, we failed to detect significant differences in GSL profiles between the genotypes (**Fig. 6C, 6D**). Using matrix-assisted laser desorption/ionization mass spectrometry (MALDI-MS) imaging, we confirmed that the major GSL (4MSOB) is increased in the sepal epidermis of *gir1* in comparison to wild type and *atml1* (**Fig. S9**).

We superimposed the results from our transcriptome analysis and GSL profiles on a current model of the GSL biosynthesis pathway from Arabidopsis (**Fig. 6E**). In this model, genes upregulated in *gir1* encode enzymes that are predicted to contribute to the increased accumulation of GSLs observed in *gir1*. Therefore, the nearly global upregulation of GSL gene expression can explain the increased levels of GSLs detected in sepals from *gir1* mutants.

## Discussion

We propose a model for the role of the ATML1-GIR1-TPL/TPR tripartite complex in the repression of GSL biosynthesis as well as the repression of giant cell elongation growth (**Fig. 7**). GIR1 interacts with ATML1 STAD through its Zn finger, and this interaction is required for GIR1 activity in the sepal. A detailed molecular understanding of the binding interface could identify novel ways to control GIR1-ATML1 interaction that in turn dictates GSL accumulation in the sepal, or in other specified tissues. It is noteworthy that the *MYB29* paralog, *MYB28*, showed an upregulated trend (**Fig. 5F**) whereas the mRNA levels of other described transcriptional regulators of the GSL pathway were not significantly altered in sepals, suggesting that the ATML1-GIR1-TPL/TPR module could play an important role in transcriptional regulation of the pathway in a tissue- or cell-type specific manner.

**Figure 7.**
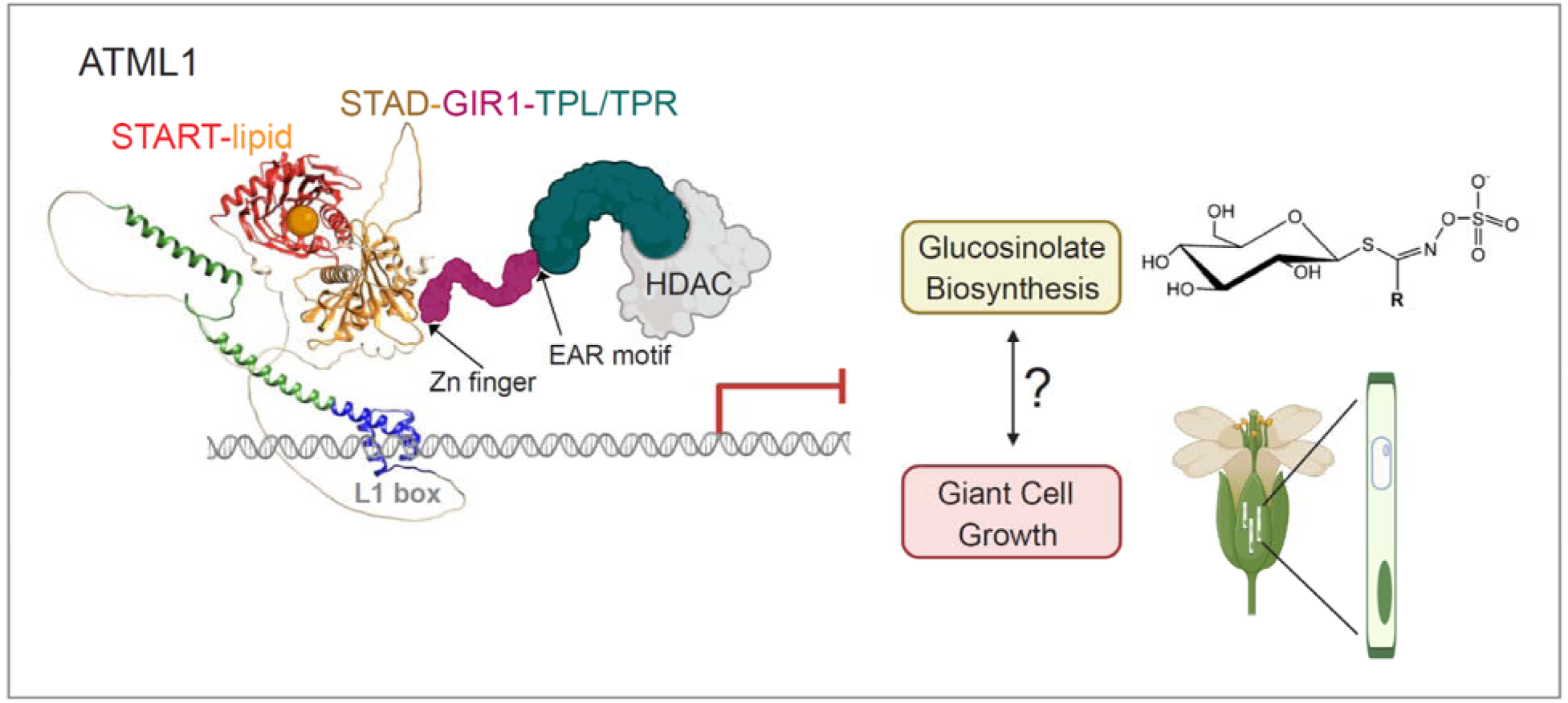
Model for biological function of ATML1-GIR1-TPL/TPR complex in repression of GSL biosynthesis and giant cell growth. The HD-Zip IV transcription factor ATML1 binds to the L1 box of target genes ^28^. The START domain of ATML1 binds a lipid ^60^, while the C-terminal STAD segment binds to GIR1 through its Zn finger. In turn, GIR1 binds to TPL/TPR corepressor proteins through its EAR motif. The TPL /TPR proteins interact with histone deacetylases (HDAC) to induce closed chromatin and transcriptional repression ^61,62^. GIR1 activity is required for both GSL biosynthesis and giant cell formation in sepals. While GIR1 appears to negatively regulate both processes, the connection between GSL biosynthesis and giant cell formation is unclear.

Mutant analysis revealed that GIR1 controls giant cell formation in sepals, but a mutant of the closely related GIR2 does not display this giant cell phenotype nor is there an additive effect in the double mutant **(Fig. 1C, Figure S1E)**. Our GSL profiling of *gir2* mutant sepals also showed no differences to wild type (**Dataset S8**). Perhaps GIR2 has a similar role to GIR1 in regulating cell elongation and/or GSL synthesis in other tissues or cell types besides sepals. This broader view is supported by work in other plant systems. ABAXIALLY CURLED LEAF1 (ACL1), a rice ortholog of GIR1, drives bulliform cell division and cell expansion, resulting in abaxial leaf curling when overexpressed ^44^. Thus, GIR1-like proteins can influence epidermal development by altering both cell proliferation and enlargement in diverse species. A recent investigation utilizing proximity labeling demonstrated that ACL1 directly interacts with the HD-Zip IV transcription factors RICE OUTERMOST CELL-SPECIFIC GENE 4 (ROC4) and ROC5, thereby negatively modulating their activities through corepressor recruitment ^45^. This is consistent with our findings that GIR1 bridges the interaction between ATML1 and TPL/TPR corepressors, consequentially resulting in decreased ATML1 target gene activation.

The *gir1* mutant displays two phenotypes specifically in the sepal epidermis: (i) excess giant cell formation and (ii) elevated GSL levels. This raises an important question about the relationship between GSLs and cell expansion. Giant cells are elongated, outward bulging cells in the epidermis and therefore they may be easily subject to insect attack. Perhaps GSLs accumulate preferentially in giant cells, providing a selective advantage to prevent insect herbivory. On the other hand, it has been reported that GSL derivatives can act as signaling molecules to influence cell growth ^46^. This is consistent with the idea that regulation of GSL accumulation may consequently control endoreduplication and cell size. There was no clear correspondence between MALDI-MS images of GSLs and the locations of the giant cells in optical images (**Fig. S9**). However, the distribution of 4MSOB appears similar to that of the flavonoid controls. One possible explanation for the non-uniform distributions of these compounds is variation in laser sampling depth due to cuticular and cell wall heterogeneity. A spatial understanding of the GSL pathway in *gir1* mutants will aid in deciphering how the GIR1-ATML1 regulatory module can be leveraged to modify GSL profiles in crops.

Alternative pathways may provide means to regulate GSL homeostasis through ATML1. A recent study implicated that leaf ER bodies accumulate β-glucosidases to hydrolyze GSLs for defense purposes and showed that *ATML1* drives this process ^47^. Therefore, ATML1-mediated regulation of leaf ER bodies could control timely response to biotic attack by shifting the balance of storage and hydrolysis of GSLs. Functionally, ATML1-regulated ER body formation may connect developmental patterning with chemical defense through the cellular localization of enzymes required for GSL bioactivation.

In addition to their role in human health ^4,5^, GSL secondary metabolites display versatile applications in agriculture. On one hand, GSLs and their derivatives possess pesticidal and antimicrobial activities ^2,3^, and therefore high levels of GSLs are advantageous in floral tissues which are especially vulnerable to herbivory and pathogen attack. On the other hand, high levels of GSLs in camelina seed meals are detrimental for cattle feeds ^48,49^. The regulatory mechanism that we have identified in Arabidopsis addresses these opposing needs, warranting the future analyses of the newly identified repression module in seeds and other relevant plant tissues.

## Methods

### Plant growth conditions and transformation

Arabidopsis plants were of the Columbia (Col-0) ecotype. T-DNA insertion lines *gir1-1* (SALK_089888C), *gir2-1* (SALK_048394) and *atml1-3* (SALK_033408) were previously described ^30,31^. The *gir1-1;atml1-3* and *gir1-1;gir1-2* double mutants were obtained by crossing. Plants were grown in BM6 (Berger, Canada) soil supplemented with vermiculite and perlite (Hummert International, Topeka, KS) at a 28:4:1 ratio. Sown seeds were stratified at 4°C for 4-6 days and transferred to 23°C under continuous light. Plants were grown in diurnal 14 h light/10 h dark conditions for RNA-seq and GSL analysis. The *gir1* plants were transformed by floral dip using *Agrobacterium tumefaciens* strain GV3101 (pMP90) ^50^. Transformants were selected on 0.8% agar medium supplemented with 1 mM KNO_3_ and 20 μg/mL Hygromycin B for native promoter constructs and 80 μg/mL kanamycin for CaMV 35S promoter constructs. At least ten transformants per line were transferred to the soil. T1 plants were screened for T2 segregation ratio of 3:1 of the relevant antibiotic resistance or EGFP expression. For each transgene, at least three to four transformants were characterized, and at least one representative homozygous line, verified by sequencing, was selected for further study.

### Construction of binary vectors and other plasmids

The cDNA sequences for *ATML1* and *GIR1* were PCR-amplified and cloned using the pENTR/D-TOPO Cloning Kit (Invitrogen, Carlsbad, CA). pENTR/D-TOPO plasmids for *gir1* mutants (Δ*EAR, R23L, R30E, R23L;R30L, C71A, C87A*) and *atml1* mutants (*K547E, F578A, D589L, N593A, N627A, E633L, K733R*) were created from wild-type templates using the Q5 Site-Directed Mutagenesis Kit (New England Biolabs). Gateway cloning with LR Clonase II (Invitrogen) was used to construct the following: 35S promoter constructs were generated by transferring pENTR/D-TOPO or pDONR207 sequences to pK7WGF2 ^51^ for EGFP fusions and pEarleyGate 201^52^ for hemagglutinin (3xHA) tag fusions; and Y2H constructs were generated by transferring pENTR/D-TOPO sequences to pDEST32 and pDEST22 plasmids (ProQuest, Invitrogen). PCR amplification with Q5 High Fidelity DNA Polymerase (New England Biolabs), gel purification with Nucleospin Gel and PCR Clean-up (Machery-Nagel), and NEBuilder HiFi DNA Assembly (New England Biolabs) into the *Hind*III and *Kpn*I linearized SR54, a derivative of pCAMBIA1300, were used to generate *GIR1* promoter constructs. For *proGIR1:GIR1*, a ∼5 kb fragment containing the *GIR1* promoter, 5’ UTR, gene body, 3’ UTR, and 340 bp downstream sequence was taken. For the *proGIR1:gir1*^Δ*EAR*^ construct, the genomic region was amplified as two segments: an upstream fragment prior to the EAR motif and a downstream fragment with the remaining sequence. For the *proGIR1:gir1^C71A^*and *proGIR1:gir1^C87A^* constructs, the genomic region was similarly amplified in two segments to incorporate the respective Cys to Ala substitution mutations. Four-fragment DNA assembly was performed to generate the *proGIR1:gir1*^Δ*EAR*^*:TPL* construct. The first segment was identical to first segment for the *proGIR1:gir1*^Δ*EAR*^ construct. The second segment contained the region downstream of the EAR motif through the last amino acid of *GIR1*. The third fragment contained the TPL C-terminal domain (944 residues). The fourth fragment contained the *GIR1* 3’-UTR and the 340 bp downstream sequence. Oligonucleotides utilized for plasmid construction are listed in **Table S2**.

### Recombinant protein production

The cDNA sequences for *GIR1* and *gir1^C71A^* were amplified and assembled into *BamH*I/*Not*I digested pT7HMT ^53^ using the NEBuilder HiFi DNA Assembly Kit (New England Biolabs). Similarly, *ATML1 STAD* and *atml1 STAD^D589L^* cDNA sequences were assembled into *Ssp*I-digested pET-His6-MBP-TEV-Lic (Addgene, Watertown, MA). Colonies from freshly transformed BL21 Rosetta (DE3) (Novagen, MilliporeSigma, Burlington, MA) were inoculated in 10 mL LB and grown at 37°C overnight. Primary cell cultures were transferred to 1 L LB and grown at 37°C to an OD_600_ of 0.8 prior to induction with 1 mM IPTG overnight at 18°C. Cells were collected by centrifugation and frozen at -80°C. Pellets were resuspended in native binding buffer (20 mM Tris [pH 8.0], 10 mM imidazole, 500 mM NaCl). Cells were lysed with a LM10 Microfluidizer Processor (Microfluidics, Westwood, MA) followed by high-speed centrifugation for 30 min. Clarified lysate was purified using a Ni affinity column and eluted in native elution buffer (20 mM Tris [pH 8.0], 500 mM imidazole, 500 mM NaCl). The proteins were further purified with a Superdex 75pg 26/600 size exclusion column in HEPES [pH 7.4]. Lyophilized MBP was a gift from Brian Geisbrecht.

### Surface plasmon resonance

ATML1 STAD, atml1 STAD^D589L^, and MBP alone were immobilized on a CMD200 sensor chip (XanTec bioanalytics) by amine coupling using a concentration of 20 μg/ml in 10 mM sodium acetate [pH 4.0]. A final immobilization density of 1138.2, 1066.5, and 1049.6 resonance units (RU) was obtained, respectively. Assays were performed in HBS-T running buffer (10 mM HEPES [pH 7.3], 140 mM NaCl, 0.005% [v/v] Tween-20) at a 30 μl/min flow rate. Analytes were diluted into running buffer before injection. Interactions were assessed in multicycle experiments with an analyte injection series (0.08, 0.16, 0.31, 0.63, 1.25, 2.5, 5, and 10 μM) applied over an association time of 120 s, followed by a dissociation time of 180 s. Surfaces were regenerated with a single 60 s injection of 2 M NaCl and 3 mM EDTA. Each injection series was fit to a steady-state 1:1 binding model (Langmuir) to calculate an equilibrium dissociation constant (*K_D_*) using Biacore T200 Evaluation Software (v 3.2, Cytiva). SD were calculated based on a triplicate injection series (n = 3).

### Y2H assays

The pDEST32 bait and pDEST22 prey vectors were transformed in opposite mating-type strains Y2HGold and Y187 as described in ^54^ using standard LiAc/PEG transformation. To assay the bait-prey interactions, mating was carried out on yeast peptone dextrose adenine (YPDA) medium to produce diploids containing both the prey and bait vectors. Following incubation for 1-2 days, diploid cells were selected on -Trp-Leu medium. Diploids were grown in liquid -Trp-Leu media to an OD600 of 1.0. From the normalized cultures, four serial dilutions were prepared and placed on permissive and selective media using a 48-pin multiplex plating tool ("Frogger," Dankar, Inc.) to assay reporter gene expression indicating interactions. Bait and prey vectors lacking inserts were used as negative controls. Confirmation of bait protein expression involved the extraction of total yeast proteins with the NaOH/trichloroacetic acid method.

### Transient protein expression and coimmunoprecipitation

*Agrobacterium* cultures harboring bait, prey, or p19 ^55^ plasmids were grown overnight. Cell mixtures with equal amounts of bait and prey plasmids (final OD600 = 0.4) were combined with cells harboring p19 (final OD600 = 0.2) and were pelleted and resuspended in 1 mL infiltration media (10 mM MES [pH 5.6], 10 mM MgCl_2_, 150 µM acetosyringone, 6 mg/ml glucose). Cells were incubated in the dark for 1 h. Leaves from 4–6-week-old *N. benthamiana* plants grown under 14 h light /10 h dark cycles were infiltrated on the abaxial side with a 1 ml needleless syringe. EGFP expression was assessed after 48 h by epifluorescence microscopy, and expressing regions were cut, weighed, flash-frozen, and stored at - 80°C. Frozen tissue (0.5 g) was ground in liquid nitrogen, and resuspended in 2.5 mL extraction buffer (150 mM NaCl, 50 mM Tris-HCl [pH 7.5], 5% [v/v] glycerol, 1 mM EDTA, 0.5% [v/v] Triton X-100, 1 mM dithiothreitol (DTT), 1X protease inhibitor (Sigma, P9599), 1 mM phenylmethanesulfonylfluoride (PMSF), 2% [w/v] polyvinylpolypyrrolidone (PVPP). Lysates were vortexed for 1 min and incubated on ice for 30 min before pelleting at 20K × *g* for 10 min at 4°C. Lysates (100 µL) were heated for 5 min at 99°C, combined with 100 µl of 2X Laemmli buffer and stored at -20°C for use as input controls. Cleared lysates were incubated with 15 µL GFP-Trap magnetic agarose beads (Chromotek) or anti-HA magnetic beads (ThermoFisher Scientific, 88836) for 60 min at 4°C with continuous end-to-end rotation. Beads were pelleted and supernatants were removed, followed by multiple 5 min washes (150 mM NaCl, 50 mM Tris [pH 7.5], 5% [v/v] glycerol, 1 mM EDTA, 0.5% [v/v] Triton X-100, 1 mM DTT, 1X protease inhibitor, 1mM PMSF) with continuous end-to-end rotation. Beads were eluted in 100 µL Laemmli buffer and resolved by SDS-PAGE followed by western blot with rabbit anti-GFP (1:2000, Invitrogen, A-11122) or rat anti-HA (1:5000, Roche, 11867431001) primary antibodies and goat anti-rabbit (1:2000; Enzo, ADI-SAB-300) or anti-rat (1:3000; Millipore, AP136P) secondary antibodies. SuperSignal West Femto Maximum Sensitivity Substrate (ThermoFisher Scientific) was used for chemiluminescent detection with an Azure c300 imager (Azure Biosystems).

### Phenotypic characterization and microscopy

Intact flower buds were dissected using fine forceps and transferred to double stick tape for examination. Images were captured with a Leica DFC295 digital camera mounted on a Leica M125 fluorescence stereo microscope. Expression of EGFP constructs was analyzed using the GFP2 filter. Scanning electron microscope (SEM) images were recorded on an SNE-Alpha high-resolution tabletop SEM with a 5-axis XYZRT stage at 5 nm resolution (Nanoimages, Lafayette, CA). Giant cell quantification was performed by marking giant cells based on characteristic morphological features followed by measuring the widths using ImageJ.

### RNA extraction and transcriptomics analysis

Stage 12 unopened flowers were dissected to obtain 200 sepals (from 50 buds) for each genotype in triplicate. Samples were frozen in liquid nitrogen prior to RNA extraction using the QIAGEN RNeasy Plant Mini Kit (Venlo, Netherlands). RNA quality evaluation, cDNA library construction, and Illumina sequencing were performed by Novogene (Sacramento, CA). Genes were identified as differentially expressed if their adjusted p value ≤ 0.05 and log2(fold change) ≥ 1. Mainstream hierarchical clustering grouped genes by their FPKM values. Gene ontology and KEGG enrichment analyses were performed for corresponding genes with an adjusted p value < 0.05.

### GSL extraction and analysis

GSL contents were measured as previously described ^56^. Immediately after tissue collection each sample was placed in 1 mL of 100% methanol. Samples were homogenized for 3 min in a paint shaker, centrifuged, and 400 μL of supernatants were transferred to a 96-well filter plate with DEAE sephadex. The filter plate was washed with water, 90% methanol and water again. Sephadex-bound GSLs were eluted after an overnight incubation with 110 μL of sulfatase. Individual desulfo-GSLs within each sample were separated and detected by HPLC-DAD, identified, and quantified by comparison to standard curves from purified desulfo-GSLs ^57^.

## Author Contributions

K.S., B.A., L.E.A., D.J.K., S.N., and Y.-J.L. designed research; L.E.A., B.A., A.U., A.A.R., S.N., S.R.L., A.K.B., A.L.W., T.R.N., B.L.G. and K.S. performed research; L.E.A., B.A., A.U, A.A.R., S.N., S.R.L, A.K.B., A.L.W., Y.-J.L., B.L.G., D.J.K. and K.S. analyzed data; and L.E.A., B.A. and K.S. wrote the paper.

## Competing Interest Statement

The authors declare no competing interest.

## Acknowledgments

We thank Adrienne Roeder for *atml1-3* seeds and Vitaly Citovsky for *gir1-1* and *gir2-1* seeds. Amie Norton and Prem Singh Thapa Chetri provided technical assistance with SEM. This work was funded by the National Science Foundation (MCB1616818; MCB2545120), National Institute of General Medical Sciences of the National Institute of Health under award no. P20GM103418, USDA National Institute of Food and Agriculture Hatch/Multi-State project 7001195, and the Johnson Cancer Research Center at Kansas State University. This is contribution no. 26-066-J from the Kansas Agricultural Experiment Station.

## Accession information

The RNA-seq raw data is available at the National Center for Biotechnology Information (NCBI) Gene Expression Omnibus (GEO) under accession GSE332896.

## Supplemental Information

Figures S1 to S9

Tables S1 to S4

Datasets S1 to S9

**Fig. S1.** The *gir1* and *gir1;gir2* mutants exhibit similarly enhanced giant cell formation.

**Fig. S2.** Overexpression of *GIR1* results in rescue of giant cell phenotype of *gir1* mutants.

**Fig. S3.** ATML1 STAD functions as a transactivation domain.

**Fig. S4.** Purification of recombinantly expressed GIR1 and ATML1 STAD wild-type and mutant proteins.

**Fig. S5.** Western blot of GIR1 proteins expressed in Y2H assays.

**Fig. S6.** Heatmap from RNA-seq data showing three biological replicates from each genotype.

**Fig. S7.** RNA-seq data for additional aliphatic GSL biosynthesis genes.

**Fig. S8.** RNA-seq data for additional indolic GSL biosynthesis genes and other GSL pathway genes.

**Fig. S9.** MALDI-MS imaging of the sepal epidermis indicates high 4MSOB GSL levels in *gir1* mutants.

**Table S1.** Steady state constants and binding affinity of GIR1 to ATML1 STAD and atml1 STAD^D589L^.

**Table S2.** Summary of expression of GSL biosynthesis genes in *gir1* sepals.

**Table S3.** Names and chemical properties of GSLs described in this study.

**Table S4.** Oligonucleotides used in this study.

**Dataset S1 (separate sheet in xls file).** Comprehensive RNA-seq data from Arabidopsis sepals.

**Dataset S2 (separate sheet in xls file).** List of DEGs that are up-regulated in *gir1* compared to wild-type sepals (padj<0.05, log2FC>1).

**Dataset S3 (separate sheet in xls file).** List of DEGs that are down-regulated in *gir1* compared to wild-type sepals (padj<0.05, log2FC>1).

**Dataset S4 (separate sheet in xls file).** List of DEGs that are up-regulated in *atml1* compared to wild-type sepals (padj<0.05, log2FC>1).

**Dataset S5 (separate sheet in xls file).** List of DEGs that are down-regulated in *atml1* compared to wild-type sepals (padj<0.05, log2FC>1).

**Dataset S6 (separate sheet in xls file).** List of DEGs that are up-regulated in *gir1* compared to *atml1* sepals (padj<0.05, log2FC>1).

**Dataset S7 (separate sheet in xls file).** List of DEGs that are down-regulated in *gir1* compared to *atml1* sepals (padj<0.05, log2FC>1).

**Dataset S8 (separate sheet in xls file).** Chemical analysis of GSL contents of mutants performed by HPLC-DAD.

**Dataset S9 (separate sheet in xls file).** Chemical analysis of GSL contents of transgenic lines performed by HPLC-DAD.

## Notes

https://www.be-md.ncbi.nlm.nih.gov/geo/query/acc.cgi?acc=GSE332896

### Competing Interest Statement

The authors have declared no competing interest.

