## Supplemental Methods, Figures and Tables for "ATML1-GIR1-TPL/TPR transcriptional repression module controls glucosinolates and giant cells in *Arabidopsis thaliana* sepals"

*Kathrin Schrick

**This PDF file includes:**

SI Methods

Figures S1 to S9

Tables S1 to S4

Legends for Datasets S1 to S9

SI References

**Other supporting materials for this manuscript include the following:**

Datasets S1 to S9

**SI Methods**

**Mass Spectrometry Imaging**

Sepals were dissected under a stereo microscope, and the abaxial side of the sepal was mounted facing up on a glass slide using a double-sided conductive carbon tape. Three sepals from each genotype were placed on the same slide to minimize batch effects. 9-aminoacridine (9-AA; 15 mg/mL in MeOH: $H_2O$ (30:70, v/v) was applied as a matrix using a TM sprayer (HTX Technologies, Chapel Hill, NC), and Au was sputter-coated for 20 s at 40 mA (Cressington 108; Ted Pella, Redding, CA) to provide surface conductivity. Matrix-assisted laser desorption/ionization mass-spectrometry imaging (MALDI-MSI) was performed using a QExactive HF Orbitrap MS (Thermo Scientific, San Jose, CA) with a MALDI source (Spectroglyph, Kennewick, WA) equipped with a 349-nm laser (Explorer One; Spectra Physics, Milpitas, CA). Data were collected in negative-ion mode for the m/z range of 400–500 with a mass resolution of 120,000 at m/z 200 to target the GSLs signals. Raster steps of 20 µm were used, considering the small size of sepals. Raw data were converted to imzML files using ImageInsight (Spectroglyph), and imzML files are loaded into the MSiReader (North Carolina State University; Raleigh, NC) software [1] to generate MS images, and METASPACE [2] was used for annotation with a false discovery rate (FDR) threshold of 10% and a mass tolerance of ± 3 ppm.

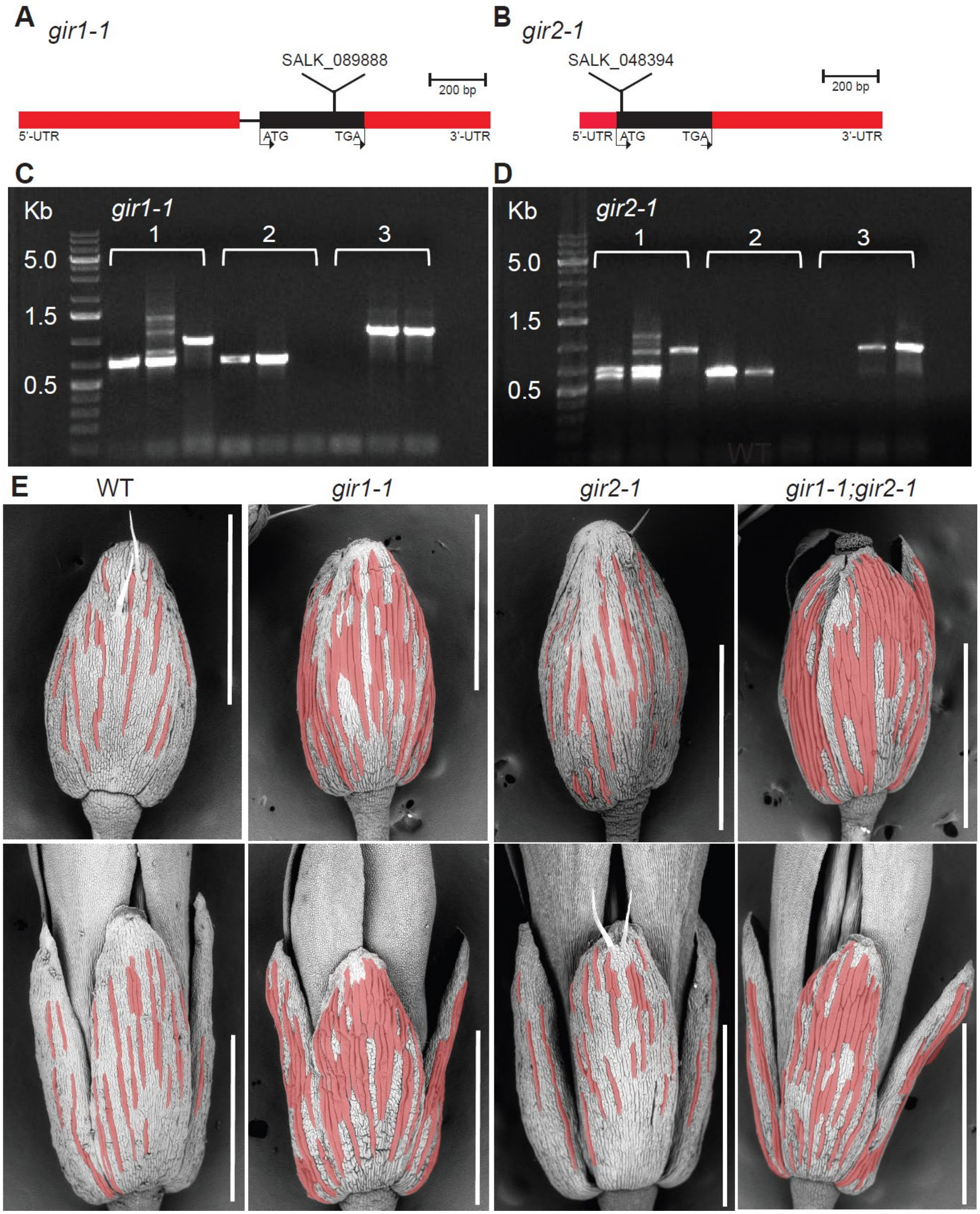


**Fig. S1. The *gir1* and *gir1;gir2* mutants exhibit similarly enhanced giant cell formation.**

**(A)** and **(B)** Schematic diagrams illustrating genomic regions encoding GIR1/At5g06270 (left) and GIR2/At3g11600 (right). The positions of the respective T-DNA insertions are indicated for *gir1-1* and *gir2-1* alleles. Untranslated regions (UTRs) are depicted in red, exons and introns are shown by black thick and thin lines, respectively.

**(C)** and **(D)** Genotyping of *gir1-1* (SALK_089888) and *gir2-1* (SALK_048394) T-DNA insertion mutants given in **(A)** and **(B)**. **Panel 1**: Amplicons from three-primer PCR with corresponding gene-specific (RP and LP) and left-border (LB) primers; **Panel 2**: Amplicons from two-primer PCR with LP and the RP gene specific primers; **Panel 3**: Amplicons from the two-primer PCR with the gene-specific primers. In each panel from left to right are shown amplicons from the mutant, heterozygote, and wild-type DNA. Oligonucleotide primers are given in **Table S4**.

**(E)** Giant cell phenotypes on sepals from wild type (WT) in comparison to *gir1, gir2,* and *gir1;gir2* mutants. Giant cell formation is similarly enhanced in *gir1 and gir1;gir2* mutants, while *gir2* mutants display a wild-type phenotype. Scanning electron microscopy (SEM) images of stage 12 (top panel) and stage 14 (bottom panel) flowers. Giant cells are falsely colored red. Bar = 1 mm. Quantification of giant cell numbers per sepal and giant cell width is given in **Fig. 1C**.

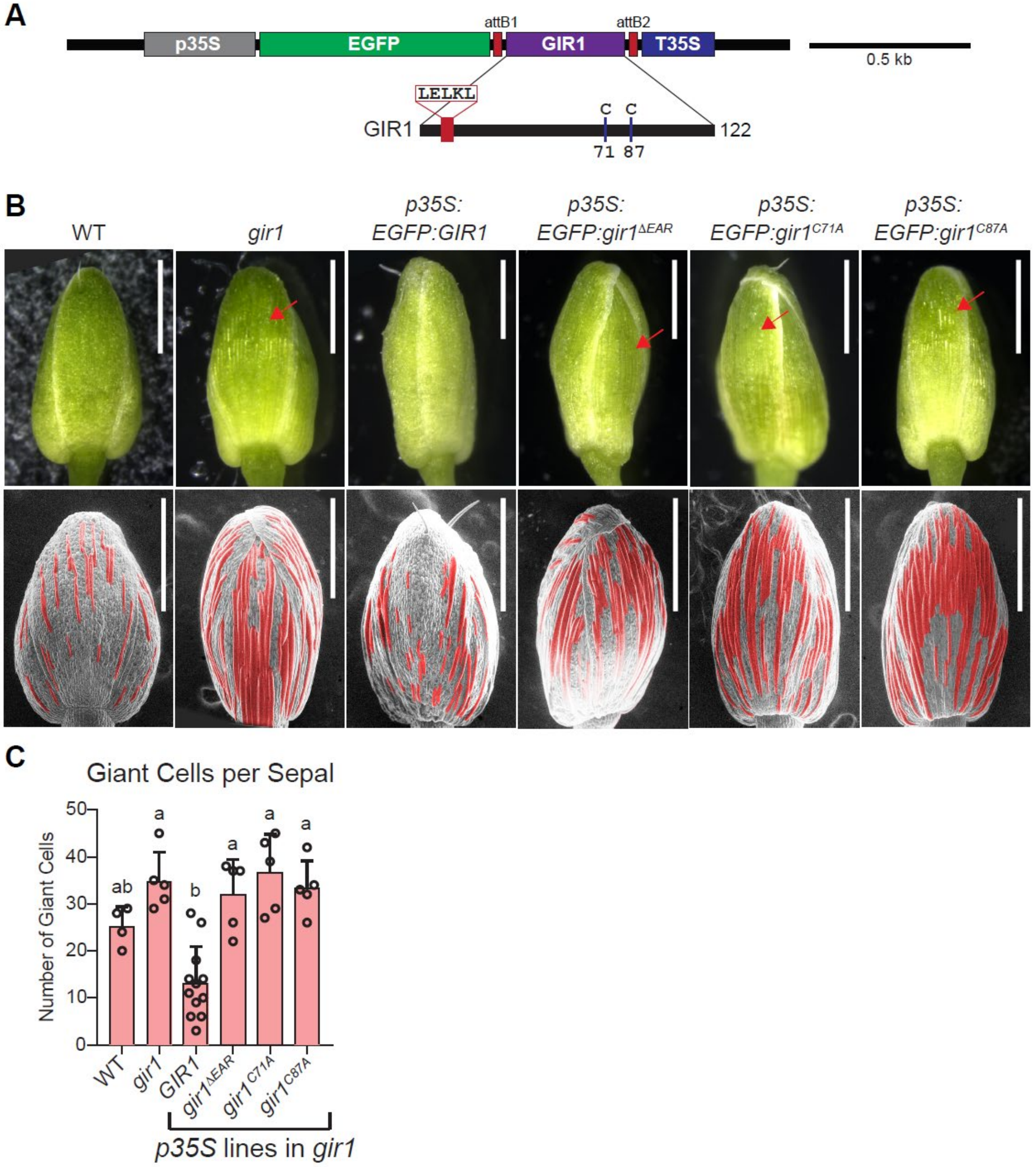


**Fig. S2. Overexpression of *GIR1* results in rescue of giant cell phenotype of *gir1* mutants.**

**(A)** Schematic of the transgenic CaMV 35S promoter (*p35S*) constructs for *GIR1* wild-type and mutants affecting the function of the EAR motif (*gir1*$^{\Delta EAR}$) and predicted Zn finger (*gir1*$^{C71A}$ and *gir1*$^{C87A}$). **(B)** Stereo microscopy images of flower buds (stage 12) (top) and matching scanning electron microscopy (SEM) images (bottom) with constructs indicated in **(A)**, in comparison to wild type and *gir1* controls. Giant cells are false-colored red. The wild-type *GIR1* construct rescued *gir1* phenotype while *gir1* mutants (*gir1*$^{\Delta EAR}$, *gir1*$^{C71A}$, *gir1*$^{C87A}$) failed to rescue. Bar = 500 μm. **(C)** Number of giant cells per sepal were quantified for the genotypes shown in **(B)**. SD are indicated for n=4-12 biological replicates. Significant differences were assessed by one-way ANOVA, Tukey's test, and are indicated by letters ($p$ value < 0.05).

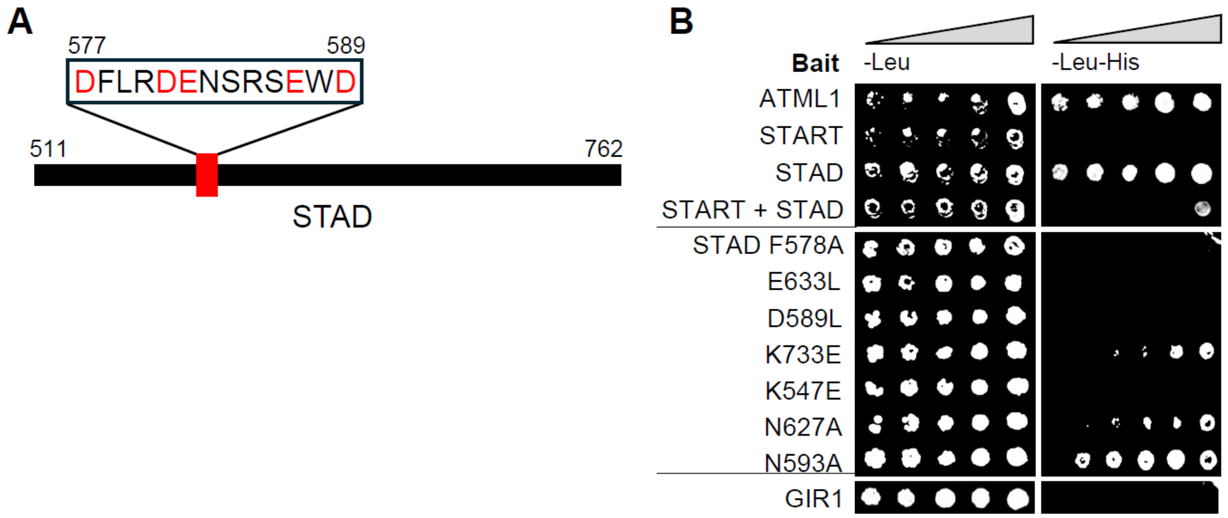


**Fig. S3. ATML1 STAD functions as a transactivation domain.**

**(A)** Schematic of ATML1 STAD is illustrated, highlighting a 13 amino acid acidic patch. Acidic amino acids, Asp (D) and Glu (E), are indicated in red. **(B)** Yeast two-hybrid (Y2H) bait autoactivation assay reveals that ATML1 STAD is an auto-activator and that specific residues in the STAD acidic patch (F578, D589) and other regions of STAD are critical for this function. Haploid yeast expressing bait plasmids with full-length ATML1, START, STAD, START+STAD, STAD mutants, or GIR1 alone were tested for transcriptional activation without prey plasmids. Growth on permissive media (-Leu) indicates presence of the bait plasmid. Growth on selective media (-Leu -His) signifies autoactivation, or transcriptional activation without requiring the prey plasmid (which contains an activation domain). Four-fold dilutions of yeast are indicated right to left.

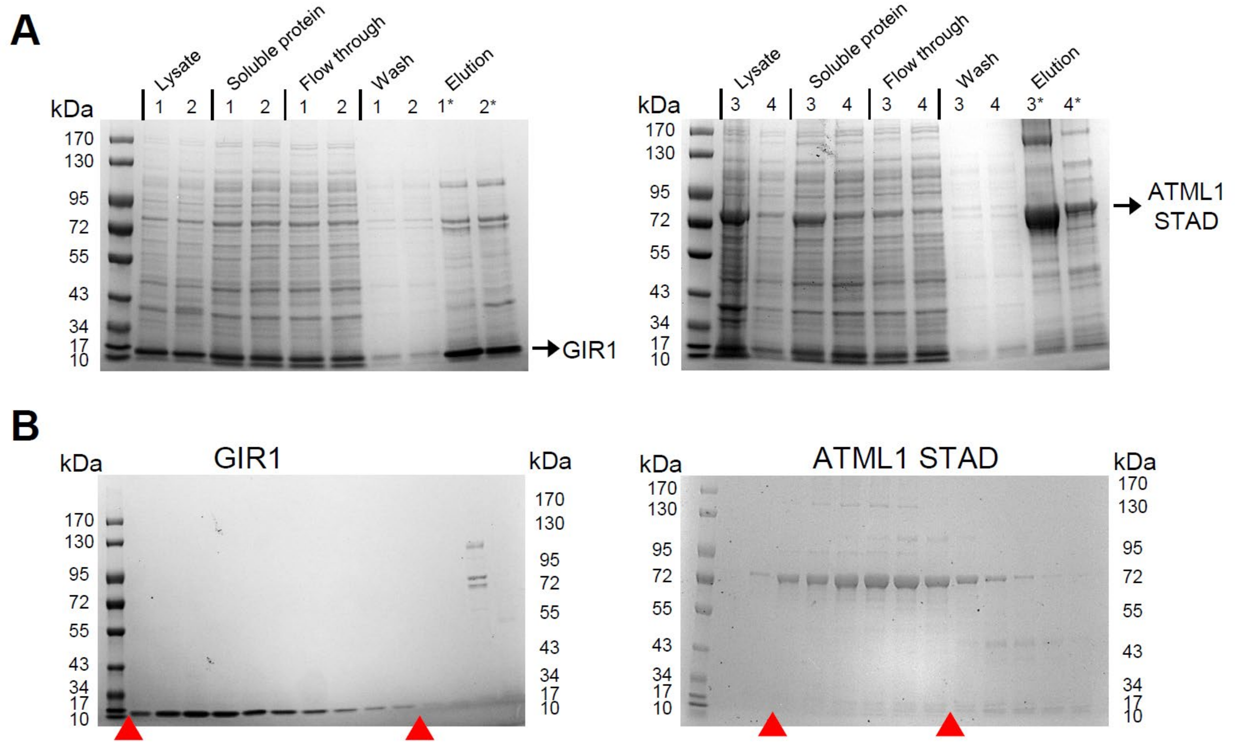


**Fig. S4. Purification of recombinantly expressed GIR1 and ATML1 STAD wild-type and mutant proteins.**

**(A)** Coomassie blue-stained gels on the left and right show protein purification of 6xHis-TEV-tagged **(1)** wild-type GIR1 and **(2)** mutant gir1$^{C71A}$, and 6xHis-MBP-TEV-tagged **(3)** wild-type ATML1 STAD and **(4)** mutant atml1 STAD$^{D589L}$, respectively. For each sample, the lysate, soluble supernatant, flow through from nickel affinity column, wash from column, and 500 mM imidazole elution are shown. Asterisks indicate samples taken for further purification by size exclusion chromatography. Arrows denote migration of the protein of interest. **(B)** SDS-PAGE of protein sample fractions purified by size exclusion chromatography. Fractions between red arrowheads indicate fractions that were pooled for final purified protein sample.

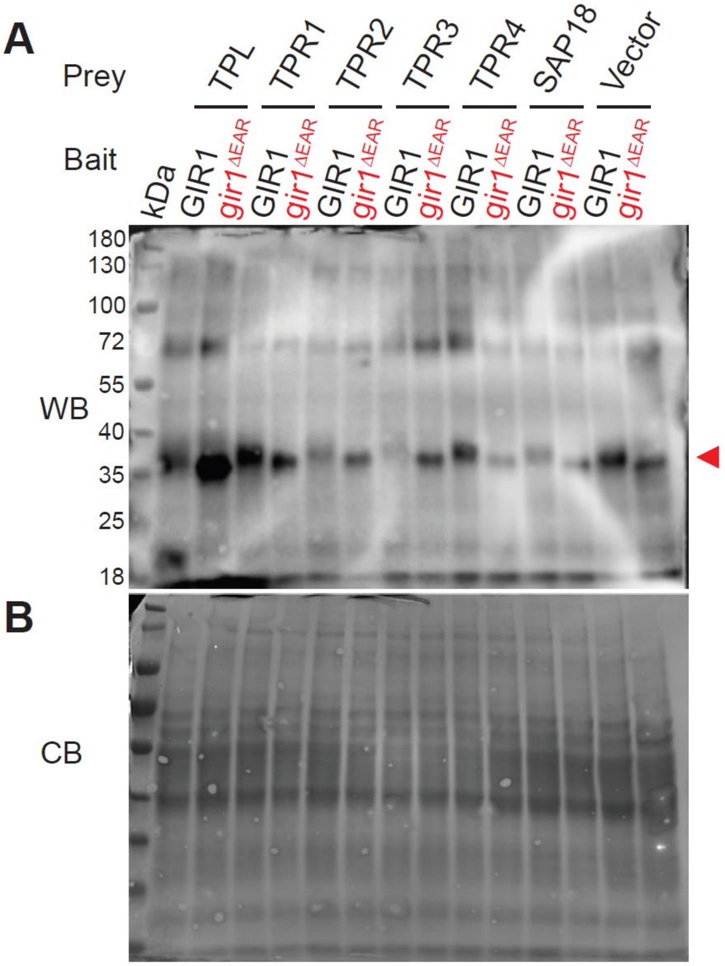


**Fig. S5. Western blot of GIR1 proteins expressed in Y2H assays.**

Western blot (WB, top) confirmed the Y2H bait protein expression for the assays shown in **Fig. 4C**. Wild-type GIR1 and mutant gir1$^{\Delta EAR}$ fused to the GAL4 DNA Binding Domain (DBD) were detected using an anti-GAL4 antibody. The red arrowhead indicates migration of GIR1 and gir1$^{\Delta EAR}$ GAL4 DBD fusion proteins. The grayscale image (bottom) shows total protein loading after Coomassie blue (CB) staining of the WB.

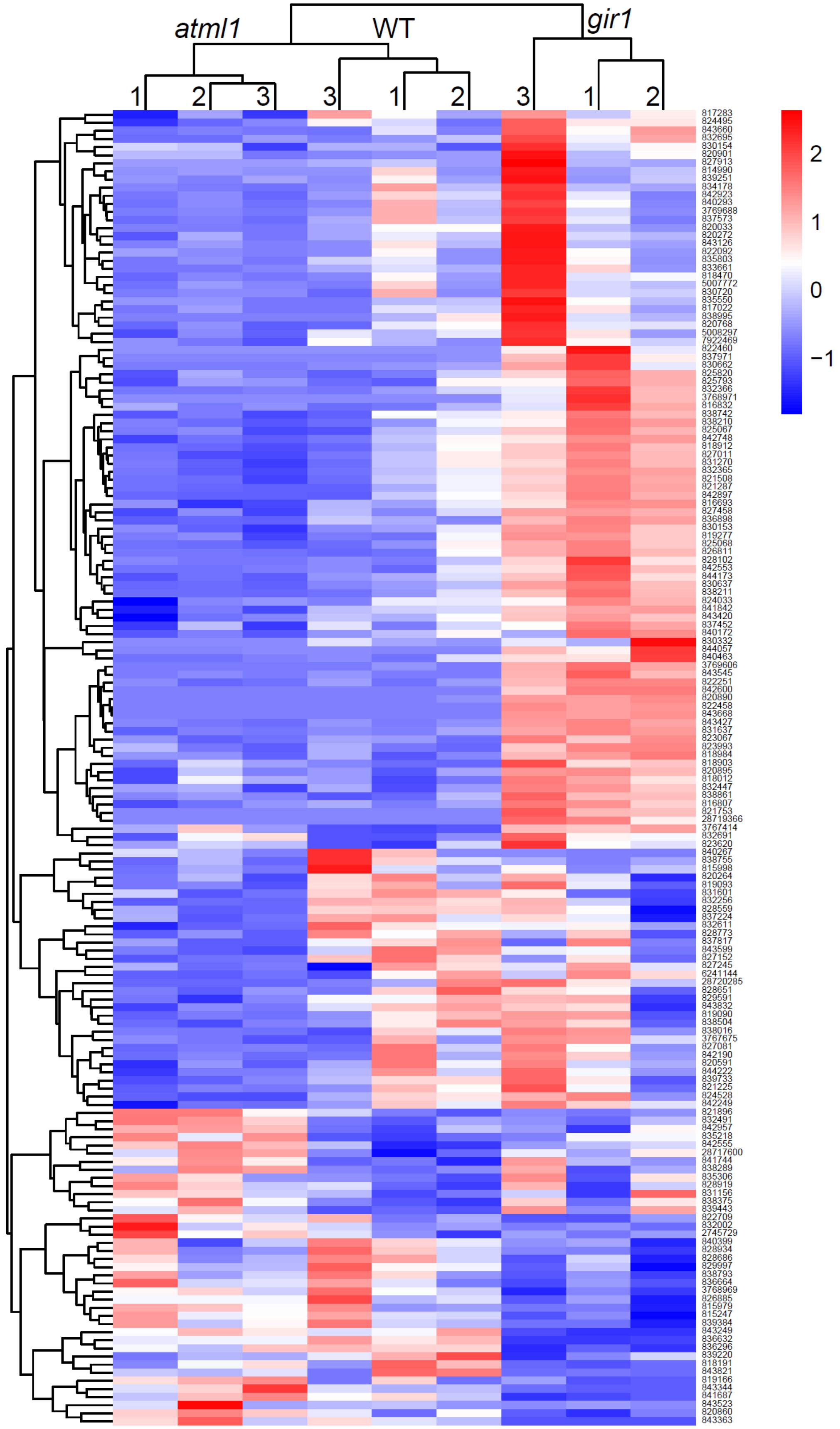

atml1
WT
gir1
1
2
3
3
1
2
3
1
2
2
1
0
−1

**Fig. S6. Heatmap from RNA-seq data showing three biological replicates for each genotype.**

Overview of DEGs in sepals from WT, *gir1*, and *atml1* plants. Gene expression levels are shown for n=3 biological replicates for each genotype, graded from low (blue) to high (red) ($P < 0.05$). A dendrogram depicting clustering of gene expression patterns is shown on the left of the heatmap. Gene IDs for the corresponding *Arabidopsis thaliana* genes are indicated on the right.

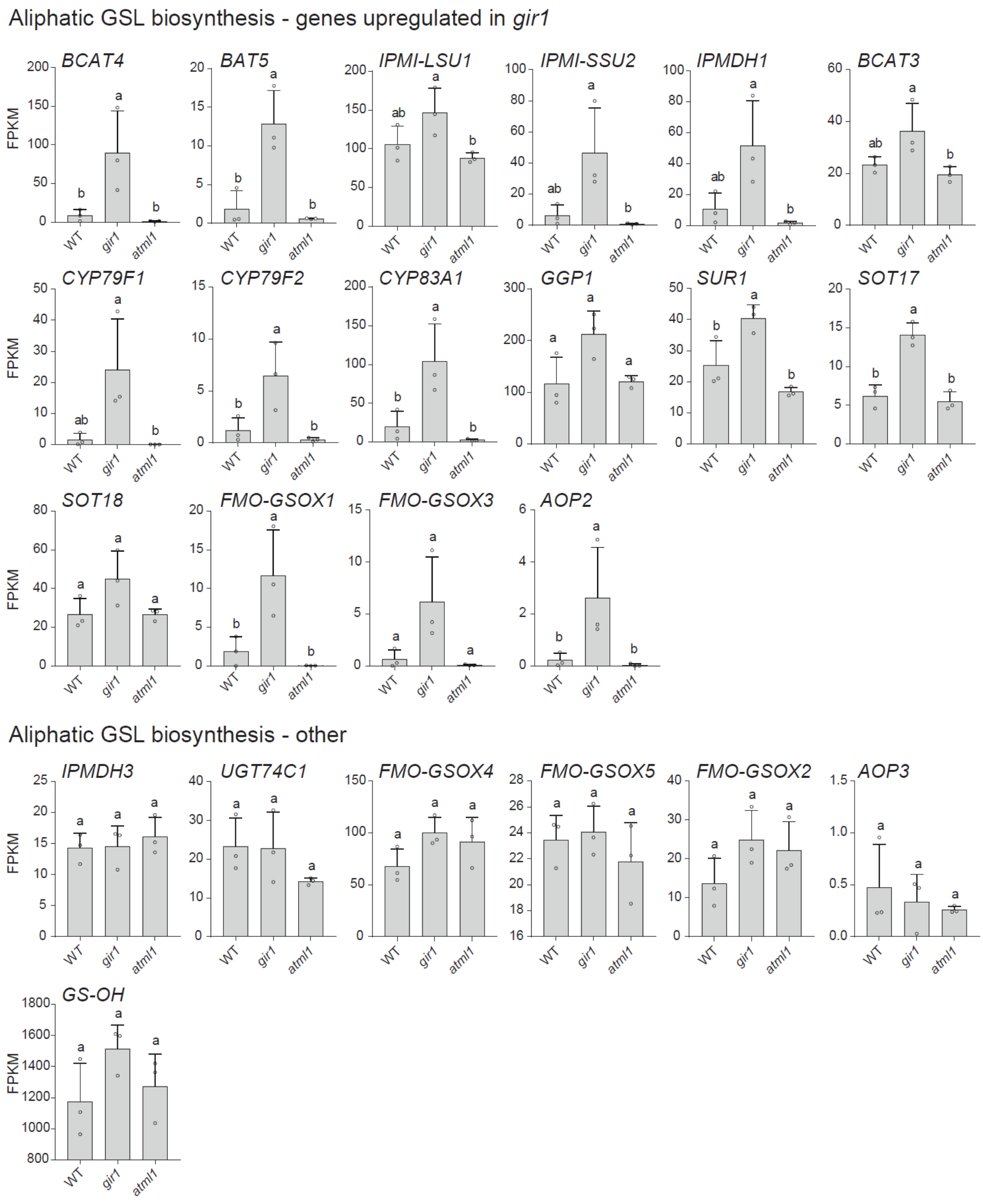


**Fig. S7. RNA-seq data for additional aliphatic GSL biosynthesis genes. Related to Fig. 5.**

The top group of graphs displays sepal RNA-seq data for aliphatic GSL biosynthesis genes upregulated in *gir1* mutant sepals, in comparison to wild type (WT) and *atml1* (*BCAT4, BAT5, IPMI-LSU1, IPMI-SSU2,*

*IPMDH1, BCAT3, CYP79F1, CYP79F2, CYP83A1, GGP1, SUR1, SOT17, SOT18, FMOGSOX1, FMOGSOX3, AOP2*). The bottom group of graphs shows sepal RNA-seq data for aliphatic GSL biosynthesis genes that are not regulated differentially in *gir1* mutants, in comparison to WT and *atml1*. (*IPMDH3, UGT74C1, FMO-GSOX4, FMOGSOX5, FMOGSOX2, AOP3, GS-OH*). FPKM, fragments per kilobase per million mapped reads. Genes are ordered within each group according to their approximate position in the GSL biosynthesis pathway, from left to right, top to bottom. Significant differences between genotypes are marked by letters (one-way ANOVA, Tukey's test, $p < 0.05$).

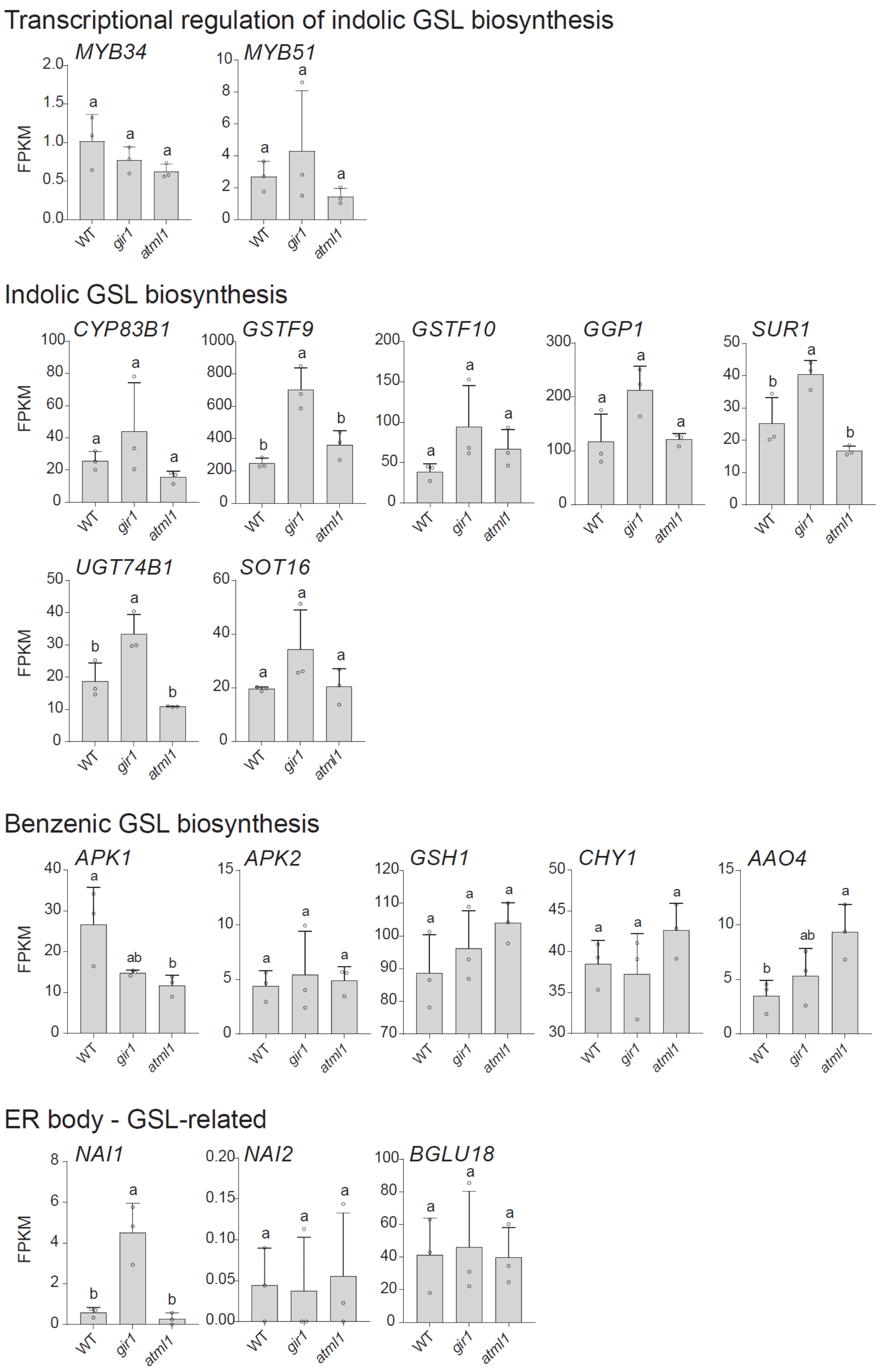


**Fig. S8. Graphs of RNA-seq data for additional indolic GSL biosynthesis genes and other GSL pathway genes. Related to Fig. 5.**

The top group of graphs displays sepal RNA-seq data for transcriptional regulators of the indolic GSL pathway (*MYB34, MYB51*). Transcripts of *MYB122* were not detected in 11/12 RNA-seq samples. The second group of graphs shows RNA-seq data for indolic GSL genes, all of which appear upregulated in *gir1* mutant sepals (*CYP83B1, GSTF9, GSTF10, GGP1, SUR1, UGT74B1, SOT16*). Note that *GGP1* and *SUR1* act in biosynthesis of both aliphatic and indolic GSLs. The third group of graphs show sepal RNA-seq data for benzenic GSL biosynthesis genes that are not upregulated in *gir1* mutants (*APK1, APK2, GSH1, CHY1, AAO4*). The bottom group of graphs show data for ER body and GSL-related genes (*NAI1, NAI2, BGLU18*). FPKM, fragments per kilobase per million mapped reads. Genes are ordered within each group according to their approximate position in the GSL biosynthesis pathway, from left to right, top to bottom. Error bars indicate SD. Significant differences between genotypes are marked by letters (one-way ANOVA, Tukey's test, $p < 0.05$).

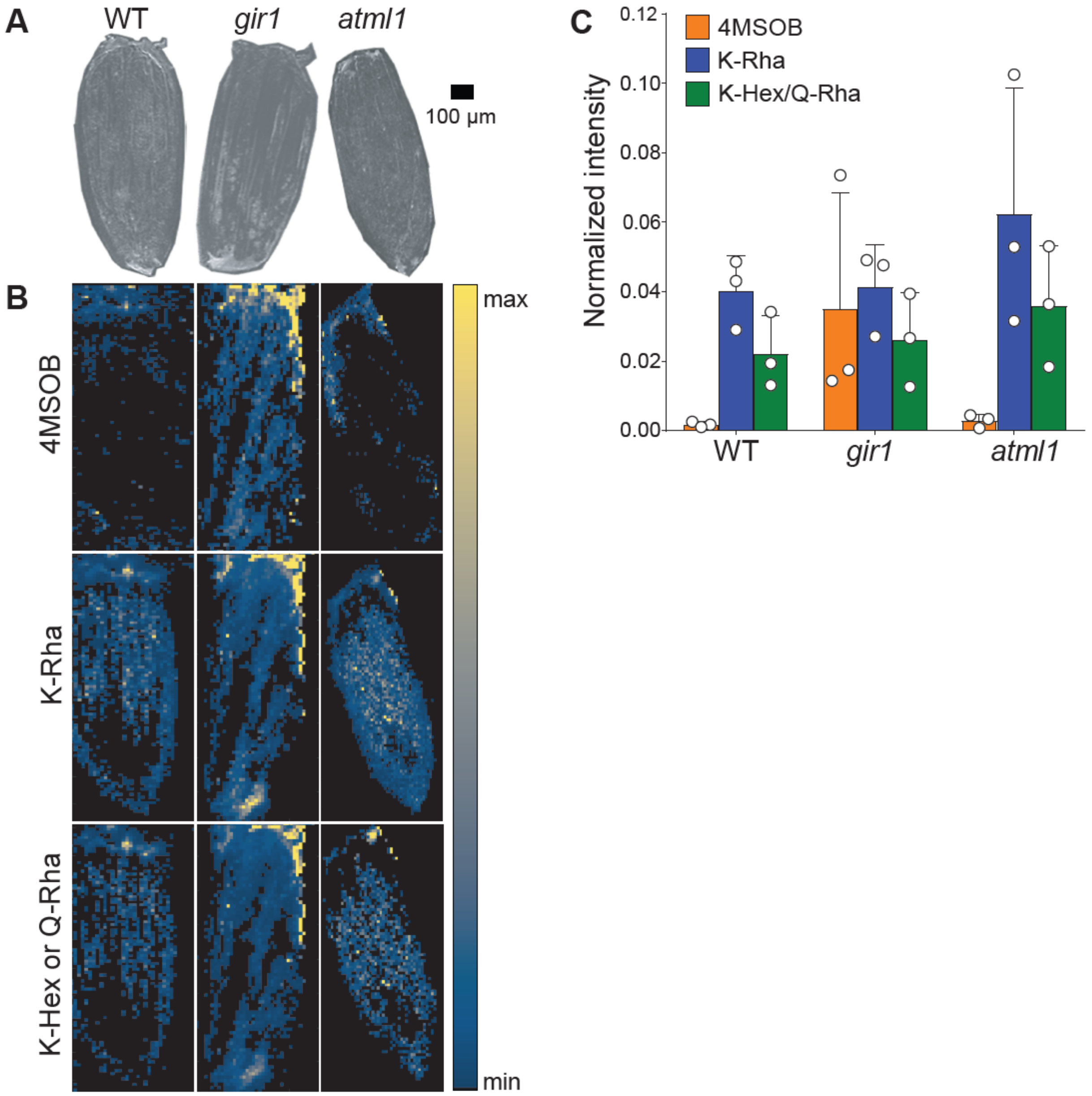


**Fig. S9. MALDI-MS imaging of the sepal epidermis indicates high 4MSOB GSL levels in the *gir1* mutants.**

**(A)** Optical images of *Arabidopsis thaliana* sepals from wild type (WT) in comparison to *gir1* and *atml1* mutants. **(B)** MALDI-MS ion images of the aliphatic GSL 4MSOB (top), and the flavonoids kaempferol rhamnoside (K-Rha) (middle), and kaempferol hexoside (K-Hex) or quercetin rhamnoside (Q-Rha) (bottom), shown to facilitate comparison with the GSL distribution. Ion intensities were normalized to the base peak (*m/z* 425.0129, $[9\text{-AA+Au+Cl–H}]^-$), and the maximum color scale was set to 1.0 across metabolites and genotypes. The color scale bar represents normalized ion intensity, and all ions are displayed as $[M–H]^-$. **(C)** Average normalized intensity of 4MSOB, K-Rha, and K-Hex or Q-Rha in each genotype at defined regions of interest (ROIs; n=3 biological replicates). Error bars indicate SD.

**Table S1. Steady state constants and binding affinity of GIR1 to ATML1 STAD and atml1 STAD$^{D589L}$.** Related to surface plasmon resonance (SPR) data presented in **Fig. 3G-H**. Equilibrium dissociation constants ($K_D$) are reported alongside the mean and SD for three injection series replicates (n=3). The data were fit to a steady-state model of binding. No fit or resulting $K_D$ could be determined for the gir1$^{C71A}$ mutant. Asterisk indicates that the $K_D$ for the GIR1 over mutant STAD$^{D589L}$ binding is significantly higher than the $K_D$ for the GIR1 over wild-type STAD binding (*t*-test, two-tailed distribution, equal variance, p-value = 0.011).

| **$K_D$ (μM)** | **GIR1 over STAD** | **GIR1 over STAD$^{D589L}$** |
|---|---|---|
| Replicate 1 | 4.08 | 7.95 |
| Replicate 2 | 4.49 | 8.24 |
| Replicate 3 | 6.04 | 9.93 |
| Mean | 4.87 | 8.71 * |
| SD | 1.03 | 1.07 |

**Table S2**. **Summary of expression of GSL biosynthesis genes in *gir1* sepals.** The fold-change (FC) difference of the average expression between the three replicates of GSL biosynthesis genes [3] in *gir1* was compared to wild type. Genes are ordered based on level of FC upregulation. Related to **Fig. 5**.

| **Expression Differences of Glucosinolate Biosynthesis Pathway Genes between *gir1* vs. WT** | | | |
|---|---|---|---|
| Gene | Gene ID | Locus | *gir1* vs. WT FC |
| **Aliphatic Pathway Genes** | | | |
| MYB29 | 830662 | At5g07690 | 16.0 |
| CYP79F1 | 838211 | At1g16410 | 14.9 |
| AOP2 | 828102 | At4g03060 | 12.0 |
| MAM1 | 832365 | At5g23010 | 10.9 |
| BCAT4 | 821508 | At3g19710 | 10.4 |
| FMO-GSOX3 | 842553 | At1g62560 | 9.5 |
| MAM3 | 832366 | At5g23020 | 7.6 |
| IPMI SSU2 | 818912 | At2g43100 | 7.3 |
| BAT5 | 826811 | At4g12030 | 6.9 |
| FMO-GSOX1 | 842897 | At1g65860 | 6.2 |
| CYP79F2 | 838210 | At1g16400 | 5.4 |
| CYP83A1 | 827011 | At4g13770 | 5.2 |
| IPMI SSU3 | 825068 | At3g58990 | 5.1 |
| IPMDH1 | 831270 | At5g14200 | 4.8 |
| GSTU20 | 844173 | At1g78370 | 4.5 |
| SOT17 | 838440 | At1g18590 | 2.3 |
| GGP1 | 829176 | At4g30530 | 1.8 |
| FMO-GSOX2 | 842551 | At1g62540 | 1.8 |
| SOT18 | 843749 | At1g74090 | 1.7 |
| BCAT3 | 824130 | At3g49680 | 1.6 |
| SUR1 | 816585 | At2g20610 | 1.6 |
| MYB28 | 836263 | At5g61420 | 1.6 |
| FMO-GSOX4 | 842554 | At1g62570 | 1.5 |
| IPMI LSU1 | 826975 | At4g13430 | 1.4 |
| GS-OH | 817083 | At2g25450 | 1.3 |
| IPMDH3 | 840006 | At1g31180 | 1.0 |
| UGT74C1 | 817736 | At2g31790 | 1.0 |
| FMO-GSOX5 | 837766 | At1g12140 | 1.0 |
| AOP3 | 828104 | At4g03050 | 0.7 |
| **Indolic and Benzenic Pathway Genes** | | | |
| CYP79B2 | 830154 | At4g39950 | 7.2 |
| CYP79B3 | 816765 | At2g22330 | 3.8 |
| GSTF9 | 817636 | At2g30860 | 2.8 |
| GSTF10 | 817637 | At2g30870 | 2.4 |
| GGP1 | 829176 | At4g30530 | 1.8 |
| UGT74B1 | 839022 | At1g24100 | 1.8 |
| SOT16 | 843750 | At1g74100 | 1.8 |
| CYP83B1 | 827011 | At4g13770 | 1.7 |
| SUR1 | 816585 | At2g20610 | 1.6 |
| **Co-substrate Pathway Genes** | | | |
| AAO4 | 839488 | At1g04580 | 1.5 |
| APK2 | 830153 | At4g39940 | 1.2 |
| GSH1 | 828409 | At4g23100 | 1.1 |
| CHY1 | 836724 | At5g65940 | 1.0 |
| APK1 | 815963 | At2g14750 | 0.6 |

**Table S3. Names and chemical properties of glucosinolates described in this study.** The glucosinolates (GSLs) are classified as aliphatic (top) or indolic (bottom).

| Abbreviation | Full Name | Common Name | Chemical Formula | Exact Mass (Da) | Molecular Weight (g/mol) |
|---|---|---|---|---|---|
| **Aliphatic GSL** | | | | | |
| 3MTP | 3-Methylthiopropyl | Glucoiberverin | C11H21NO9S3 | 407.03784477 | 407.5 |
| 4MTB | 4-Methylthiobutyl | Glucoerucin | C12H23NO9S3 | 421.05349483 | 421.5 |
| 5MTP | 5-Methylthiopentyl | Glucoberteroin | C13H25NO9S3 | 435.0691 | 435.5 |
| 6MTH | 6-Methylthiohexyl | Glucolesquerellin | C14H27NO9S3 | 449.0848 | 449.6 |
| 7MTH | 7-Methylthioheptyl | | C15H29NO9S3 | 463.10044502 | 463.6 |
| 8MTO | 8-Methylthiooctyl | | C16H31NO9S3 | 477.11609509 | 477.6 |
| 3MSOP | 3-Methylsulfinylpropyl | Glucoiberin | C11H21NO10S3 | 423.0328 | 423.5 |
| 4MSOB | 4-Methylsulfinylbutyl | Glucoraphanin | C12H23NO10S3 | 437.0484 | 437.5 |
| 5MSOP | 5-methylsulfinylpentyl | Glucoalyssin | C13H25NO10S3 | 451.0641 | 451.5 |
| 6MSOH | 6-Methylsulfinylhexyl | Glucohesperin | C14H27NO10S3 | 465.07970958 | 465.6 |
| 7MSOH | 7-Methylsulfinylheptyl | | C15H28NO10S3 | 478.08753461 | 478.6 |
| 8MSOO | 8-Methylsulfinyloctyl | Glucohirsutin | C16H31NO10S3 | 493.11100971 | 493.6 |
| 3OHP | 3-Hydroxypropyl | | C10H19NO10S2 | 377.04503815 | 377.4 |
| 4OHB | 4-hydroxybutyl | | C11H21NO10S2 | 391.06068821 | 391.4 |
| Benzyl | Benzyl | Glucotropaeolin | C14H19NO9S2 | 409.05012353 | 409.4 |
| 3BZOP | 3-Benzoyloxypropyl | Glucomalcomiin | C17H23NO11S2 | 481.07125290 | 481.5 |
| 4BZOB | 4-Benzoyloxybutyl | | C18H25NO10S2 | 479.09198834 | 479.5 |
| **Indolic GSL** | | | | | |
| I3M | Indol-3-ylmethyl | Glucobrassicin | C16H20N2O9S2 | 448.06102257 | 448.5 |
| 1OHI3M | 1-hydroxy-indol-3-ylmethyl | | C16H20N2O10S2 | 464.0559 | 464.5 |
| 4OHI3M | 4-hydroxy-indol-3-ylmethyl | | C16H19N2O10S2 | 463.04811215 | 463.5 |
| 1MOI3M | 1-methoxy-indol-3-ylmethyl | | C17H22N2O10S2 | 478.07158725 | 478.5 |
| 4MOI3M | 4-methoxy-indol-3-ylmethyl | | C17H22N2O10S2 | 478.07158725 | 478.5 |

**Table S4**. **Oligonucleotides used in this study.**

| | |
|---|---|
| Primers for genotyping mutants. | |
| gir1_SALK_089888_F | CTTTTTCTGGGGATCAGGATC |
| gir1_SALK_089888_R | AAGGCCTAAGCTCAAGCAATC |
| gir2_SALK_048394_F | CAACAAAATGGAATAACAGTTAAAGC |
| gir2_SALK_048394_R | CAGGTCTTCTTCTTGCGTCTG |
| atml1-3_F | CAGGCAGAAGAAAATCGAGAT |
| atml1-3_R | GAAACCAGTGTGGCTATTGTT |
| LbB1.3 (T-DNA for *gir1, 2*) | ATTTTGCCGATTTCGGAAC |
| LBb1 (T-DNA for *atml1-3*) | GCGTGGACCGCTTGCTGCAACT |
| Primers for TOPO directional cloning. Additional CACC sites in forward primers required for directional cloning are bolded while stop codons are in blue. Alternative bases used for mutant generation are red. | |
| Topo_ATML1_F | **CACC** ATG TAT CAT CCA AAC ATG TTC G |
| ATML1_C_R_new | AT CGA TTA GGC TCC GTC GCA G |
| ATML1_START_F | **CACC** ATA CCT TCT GAG GCT GAT AAG |
| ATML1_START_R | CTA CAT GGA ACT GGC GAG CCG |
| ATML1_STAD_F | **CACC** GCC AGC AAC ATT CCG G |
| GIR1_cDNA_F | **CACC** ATG AGT CGA AGA AGT CC |
| GIR1_cDNA_R | TCA TCA GTT CCT TCG AGT C |
| TPL_F | **CACC** ATG TCT TCT CTT AGT AG |
| TPL_R | TCA TCT CTG AGG CTG ATC AG |
| Topo_TPR1_F | **CACC** ATG TCT TCT CTG AGC AG |
| Topo_TPR1_R | TCA TCT CTG AGG CTG GTC AG |
| atml1_K547E_F | CCG GGA AGA CCT CCA GGC ATC G |
| atml1_K547E_R | ATC ATC CAT GCT CTC TCG GGT CAT GAC C |
| atml1_D589L_F | CTT ATA CTT TCC AAT GGA GGC |
| atml1_D589L_R | CCA CTC GCT TCT TGA GTT TTC ATC |
| atml1_N593A_F | GCT GGA GGC TTG GTT CAA GAA ATG |
| atml1_N593A_R | GGA AAG TAT ATC CCA CTC GCT TC |
| atml1_N627A_F | GCC ATG TTG ATC TTA CAA GAA AGT TG |
| atml1_N627A_R | GCT CTG CCC TGA GTT CCC |
| atml1_E633L_F | TTA AGT TGT ACG GAC GCA TCA GGG |
| atml1_E633L_R | TTG TAA GAT CAA CAT GTT GCT CTG CCC |
| atml1_K733E_F | GAA CTC TCT CTC GGT TCA GTT GC |
| atml1_K733E_R | AGC GGT AGG AAC AGA GTC AAC AAG |
| gir1_R23L;R30L_F | TTG ATG GTT CGA TCT CCA AGC CTC TCA GCG |
| gir1_R23L_R | CCT CTG GCT TGA CGT AGG CGG CG |
| gir1_R30E_F | GAG TCA GCG ACG ACT TCA CCG ACC |
| gir1_R30L_R | GCT TGG AGA TCG AAC CAT CCT CCT CTG G |
| Primers for DNA assembly to clone into SR54 vector. Sequence overlapping with SR54 colored in blue while restriction sites are indicated in red. Bases used for alanine mutations are bolded. | |
| SR54_gir1_complement_F | AACGACGGCCAGTGCC AAGCTT GATGAAATGCATGAACGTACG |
| SR54_gir1_complement_R | CACCGCGGTGGAGCTC GGTACC GTTTTTAGGCCTGTGATTAGC |
| gir1_C87A_F | **GC**C CCT AAA TGC AAA AGC ACC G |
| gir1_C87A_R_Ass | GT GCT TTT GCA TTT AGG G**GC** TTT AGG GTC GTC |
| gir1_C71A_F | **GC**T CCT CGT TGT CTC ATG TAC GTT ATG C |
| gir1_C71A_R_Ass | TACATGAGACAACGAGG A**GC** TCC GAC AAG AAC C |
| gir1_Del_EAR_F | AATCTCTCGCCGCCTACG |
| gir1_Del_EAR_R | CTT TGG ACT TCT TCG ACT CAT |
| gir1_Del_21_18_EAR_F | ATGAGTCGAAGAAGTCCAAA GAA TCT CTC GCC GCC TAC G |
| GIR1_TPL_R | C TGG ATT TGG CCT TGG GTT CCT TCG AGT CTT GCG AC |
| TPL_C_Term_F | CCA AGG CCA AAT CCA GAT ATC |
| TPL_R | TCA TCT CTG AGG CTG ATC AG |
| TPL_C_GIR1_3UTR_F | CAGCCTCAGAGATGATGAAGGACCAGTAAAATCAAAATCCTG |

| Primers for cloning cDNAs into plasmids for protein production. (Restriction sites, red; cDNA sequences, blue) | |
|---|---|
| pET-His6-MBP-TEV-Lic_SspI_ATML1_STAD_F | ACCTGTACTTCCAATCC AATATT AGTCCTGAGGGGAGAAA |
| pET-His6-MBP-TEV-Lic_SspI_ATML1_STAD_R | ATCCGTTATCCACTTCC AATATT TTAGGCTCCGTCGCA |
| pT7HMT_BamHI_GIR1_F | TACTTCCAGGGGTCGACA GGATCC ATGAGTCGAAGAAGTCC |
| pT7HMT_NotI_GIR1_R | GTGGTGGTGCTCGAGT GCGGCCGC TCAGTTCCTTCGAGTCTTG |

### SI Dataset Legends

**Dataset S1 (separate sheet in xls file).** Comprehensive RNA-seq data from Arabidopsis sepals. Raw read counts, followed by fpkm (fragments per kilobase of transcript for million mapped reads) normalized read counts are show for each of the three replicates for wild type (WT), *gir1* and *atml1* sepal RNA. Log2 fold-change values, p values, and adjusted p values (padj) are indicated for each binary genotype comparison. The following gene metrics for each gene ID are given:  gene name, chromosomal location (chr), start position, end position, strand (+ or -), length, biotype, gene description, gene family.

**Dataset S2 (separate sheet in xls file).** List of DEGs that are up-regulated in *gir1* compared to wild-type sepals (padj<0.05, log2FC>1).

**Dataset S3 (separate sheet in xls file).** List of DEGs that are down-regulated in *gir1* compared to wild-type sepals (padj<0.05, log2FC>1).

**Dataset S4 (separate sheet in xls file).** List of DEGs that are up-regulated in *atml1* compared to wild-type sepals (padj<0.05, log2FC>1).

**Dataset S5 (separate sheet in xls file).** List of DEGs that are down-regulated in *atml1* compared to wild-type sepals (padj<0.05, log2FC>1).

**Dataset S6 (separate sheet in xls file).** List of DEGs that are up-regulated in *gir1* compared to *atml1* sepals (padj<0.05, log2FC>1).

**Dataset S7 (separate sheet in xls file).** List of DEGs that are down-regulated in *gir1* compared to *atml1* sepals (padj<0.05, log2FC>1).

**Dataset S8 (separate sheet in xls file).** Chemical analysis of GSL contents of mutants performed by HPLC-DAD. Quantification of GSL is shown for extracts from gir*1*, *gir2*, and *atml1* mutant sepals and buds without (w/o) sepals in comparison to wild type (WT). Full names of GSL compounds and information on their properties are given in **Table S3**. GSLs were quantified from absorbance peaks.

**Dataset S9 (separate sheet in xls file).** Chemical analysis of GSL contents of transgenic lines performed by HPLC-DAD. Quantification of GSL is shown for extracts from transgenic *gir1* mutant lines carrying stable wild-type *pGIR1:GIR1* and mutant *pGIR:gir1*$^{C71A}$ transgenes. Extracts from sepals versus buds without (w/o) sepals are listed. Full names of GSL compounds and information on their properties are given in **Table S3**. GSLs were quantified from absorbance peaks.
